## Supplementary figures and tables for "PHOSPHATE STARVATION RESPONSE enables arbuscular mycorrhiza symbiosis"

**List of Supplementary Figures**

**Fig. S1.** Brightfield images of roots stained with acid-ink to visualize colonization of wild type (cv. Nipponbare), *phr2* and *35S:PHR2* and wild-type (cv. ZH11) and *phr2*(C) roots by *Rhizophagus irregularis* at 7 wpi when grown at LP (25 µM $P_i$) in quartz sand.

**Fig. S2.** Brightfield images of roots stained with acid-ink to visualize colonization of wild type (cv. Nipponbare), *phr2* and *35S:PHR2* roots by *Rhizophagus irregularis* at 7 wpi and grown at HP (500 µm $P_i$) in quartz sand.

**Fig. S3.** Effect of low (LP, 25 µM $P_i$) and medium (MP, 200 µM $P_i$) phosphate conditions on total root colonization, arbuscules and vesicles at 7 wpi.

**Fig. S4.** Number of assigned fragments in RNA-Seq along with fragments consistently mapped to the transcriptome reference.

**Fig. S5.** PCA plot for the RNA-Seq based transcriptome of mock and AMF-inoculated wild-type and *35S:PHR2* roots grown at HP conditions.

**Fig. S6.** Hierarchical clustering of combined DEGs (AM vs Mock samples, $log_2$(Fold-change), lfc) from roots of *phr2* and *35S:PHR2* in LP and HP respectively and wild type at both $P_i$ conditions.

**Fig. S7. Overlap of genes with decreased transcript levels in *phr2* in Mock or AM roots with AM genelist.**

**Fig. S8.** Expression of genes required for or induced during AM depend on PHR2.

**Fig. S9.** RT-qPCR-based transcript accumulation of selected DEGs recapitulates the RNA-Seq results.

**Fig. S10.** RT-qPCR-based transcript accumulation of selected DEGs at medium phosphate.

**Fig. S11.** ChIP-Seq binding peaks of PHR2-FLAG are enriched near the transcriptional start site.

**Fig. S12.** Motifs over-represented in DNA sequences with PHR2 binding sites.

**Fig. S13.** Binding site analysis for rice PHR2.

**Fig. S14.** IGV browser view of ChIP-Seq peaks adjacent to previously known PHR2 target genes.

**Fig. S15. Enrichment of PHR2 direct targets in AM genelist.**

**Fig. S16.** IGV browser view of ChIP-Seq peaks adjacent to AM-relevant genes**.**

**Fig. S17.** Enrichment of PHR2 at P1BS promoter motifs detected by ChIP-qPCR.

**Fig. S18. GA-related genes in RNASeq and ChIP-Seq.**

**Fig. S19. *Lotus japonicus* PHR1A protein.**

**Fig. S20. Position of P1BS motifs in the promoters of strigolactone biosynthesis genes in *Lotus japonicus*.**

**Fig. S21.** Temperature, sunlight and relative humidity profiles during the greenhouse experiment in field soil.

**Fig. S22.** PHR2 affects rice agronomic traits in a field soil.

**Fig. S23.** Root colonization and RT-qPCR-based transcript accumulation of AM-marker genes in roots of plants grown in field soil.

**Fig. S24.** Brightfield images of roots stained with acid-ink to visualize colonization of wild type (cv. Nipponbare), *phr2* and *35S:PHR2* roots by *R. irregularis* at 110 days post transplantation and grown at LP (unfertilized) or HP (fertilized with superphosphate fertilizer, $P_2O_5$) in field soil.

**Fig. S25.** RT-qPCR-based transcript accumulation of phosphate transporters and starvation marker genes in roots of plants grown in field soil.

**List of Supplementary Tables**

**Table S1.** Primers used for RT-qPCR and genotyping.

**Table S2.** Primers used for ChIP qPCR.

**Table S3.** Primers used for cloning.

**Table S4.** Plasmid construction by Golden Gate cloning (Level I, II and III).

**List of Supplementary Datasets (Available online)**

**Data S1:** RNA-seq based expression Data for all genes expressed in Mock (not inoculated) or AM (Rhizophagus irregularis-inoculated) roots of WT, *phr2* and *35S:PHR2*.

**Data S2:** Information on clustering analysis for the genes significantly regulated in the Mock roots of *phr2* vs WT grown at LP and *35S:PHR2* vs WT grown at HP samples in RNASeq.

**Data S3:** List of DEGs upregulated in wild type AM vs Mock LP and DEGs downregulated in *phr2* vs WT LP Mock roots.

**Data S4:** List of AM genes (AM genelist).

**Data S5:** Clustering analysis for the combined unique list of DEGs (14333 DEGs) significantly regulated in AM vs Mock samples of *phr2* under low phosphate (LP), *35S:PHR2* under high phosphate (HP) and WT under both phosphate conditions.

**Data S6:** Genes with reduced expression in *phr2* vs WT in AM and/or Mock samples at LP and required for or involved in AM symbiosis as determined by mutant analysis (red bold), RNAi (green) or overexpression (blue).

**Data S7:** Venn intersection of DEGs downregulated in *phr2* vs WT in AM and/or Mock samples at LP with DEGs upregulated in *smax1* or *d3/smax1* vs WT (Choi et al., 2020) and with AM genelist (Data S6).

**Data S8:** List of genes annotated to PHR2 binding sites (detected as MACS2 narrow peaks) and fasta sequence for PHR2 binding sites in ChIP-Seq of PHR2-FLAG.

**Data S9:** Occurrence analysis for P1BS/P1BS-like motifs in 3000 bp upstream sequence for genes with reduced expression in *phr2* and PHR2 targets in the two ChIP biological replicates.

**Data S10:** Direct PHR2 targets as identified by ChIP-Seq and RNASeq assays used in this work, therefore Set A + B genes in Fig. 3A and ChIP binding statistics for known targets of PHR2.

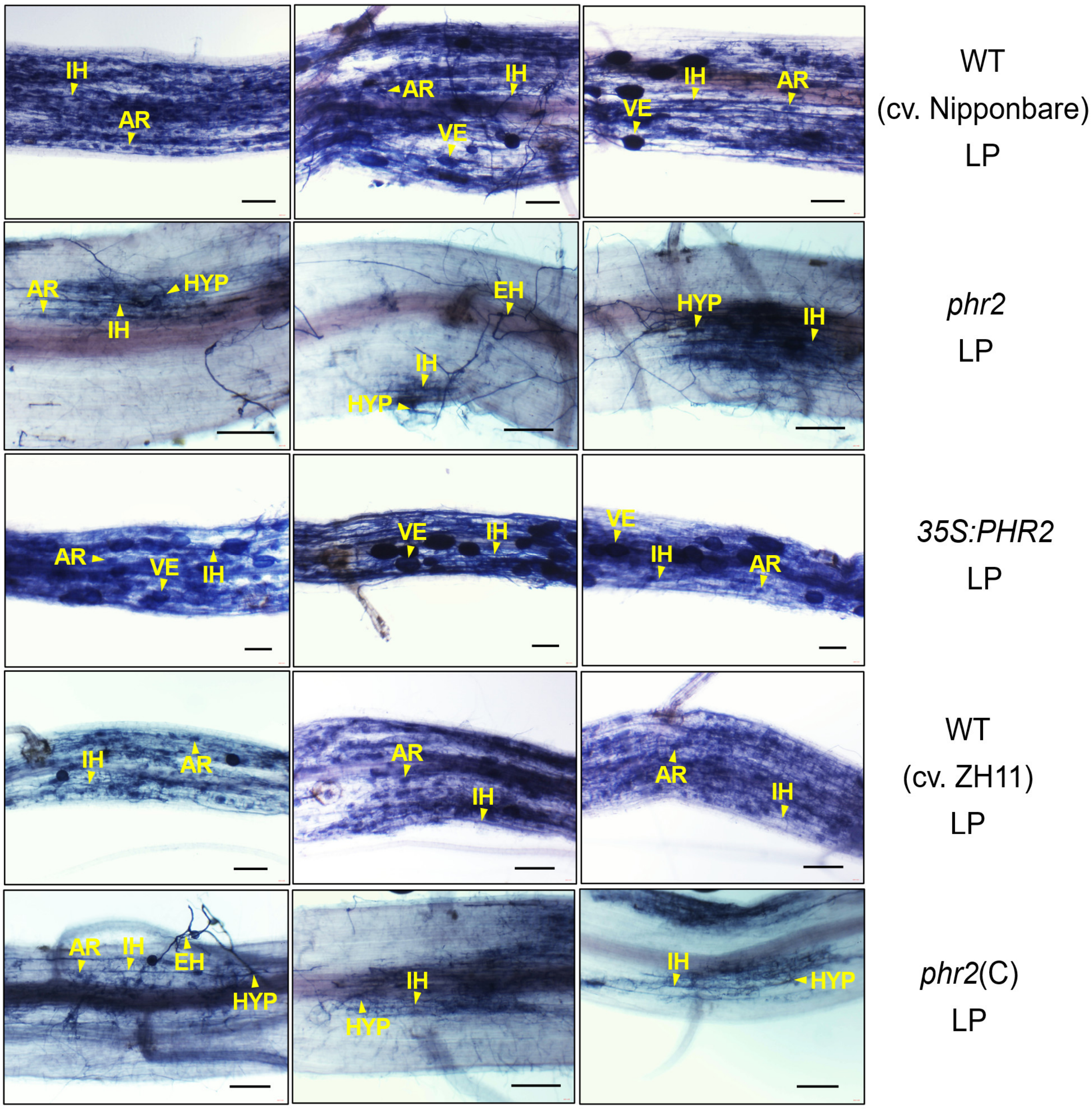

**Fig. S1.** Brightfield images of roots stained with acid-ink to visualize colonization of wild type (cv. Nipponbare), *phr2* and *35S:PHR2* and wild-type (cv. ZH11) and *phr2*(C) roots by *Rhizophagus irregularis* at 7 wpi when grown at LP (25 μM $P_i$) in quartz sand. Scale bars, 200 μm. Abbreviations: EH, extraradical hypha; HYP: hyphopodium; IH, intraradical hypha; AR, arbuscule, VE, vesicle.

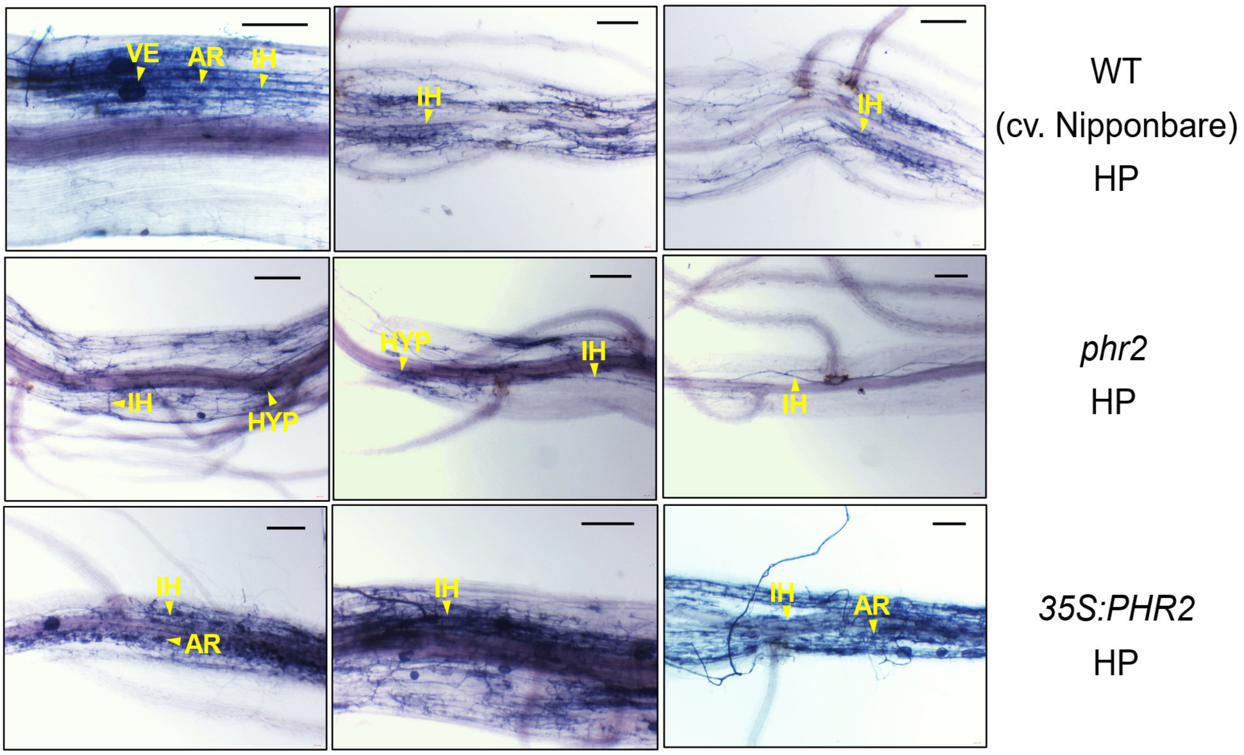

**Fig. S2.** Brightfield images of roots stained with acid-ink to visualize colonization of wild type (cv. Nipponbare), *phr2* and *35S:PHR2* roots by *Rhizophagus irregularis* at 7 wpi and grown at HP (500 µm $P_i$) in quartz sand. Scale bars, 200 µm. Abbreviations: EH, extraradical hypha; HYP: hyphopodium; IH, intraradical hypha; AR, arbuscule, VE, vesicle.

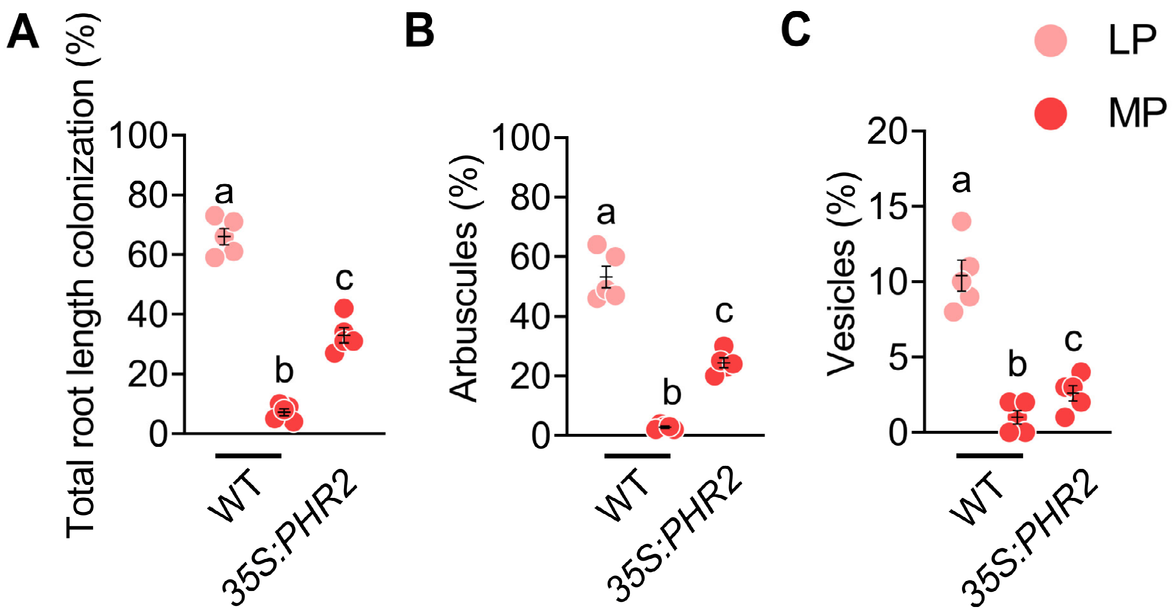

**Fig. S3.** Effect of low (LP, 25 µM $P_i$) and medium (MP, 200 µM $P_i$) phosphate conditions on total root colonization (**A**), arbuscules (**B**) and vesicles (**C**) at 7 wpi.
Statistics: Individual data-points and mean ± SE are shown. N=5; Brown-forsythe and Welch's One-Way ANOVA test with Games-Howell's multiple comparisons test. Different letters indicate statistical differences between samples.

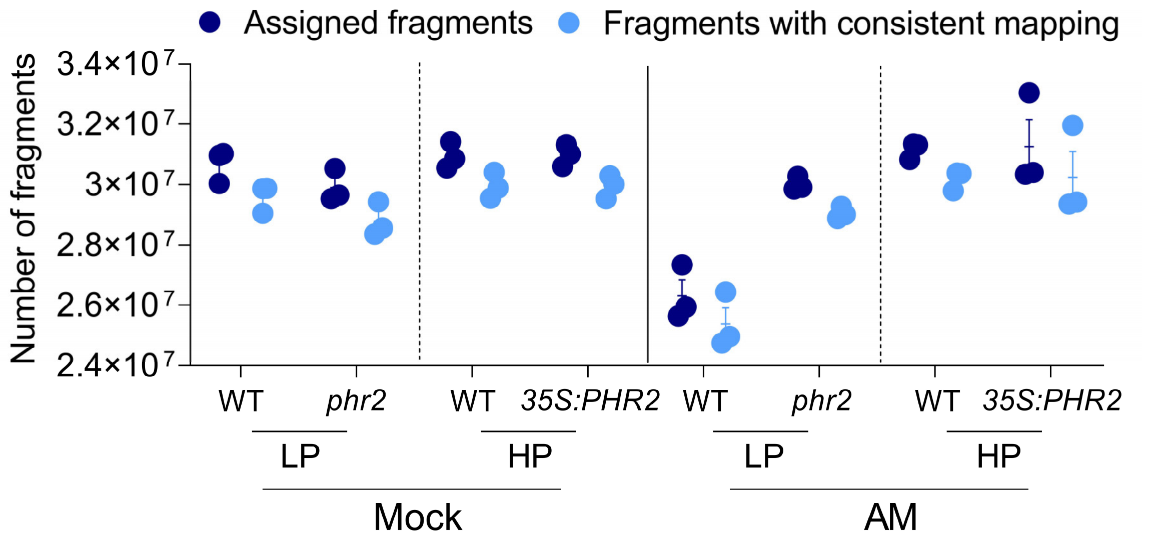

**Fig. S4.** Number of assigned fragments in RNA-Seq along with fragments consistently mapped to the transcriptome reference, n=3. Individual data-points and mean ± SE are shown. Lower number of fragments in AM-inoculated wild-type roots grown at LP results from the fact that these sample also contain reads from *R. irregularis* (≈10% of total fragments).

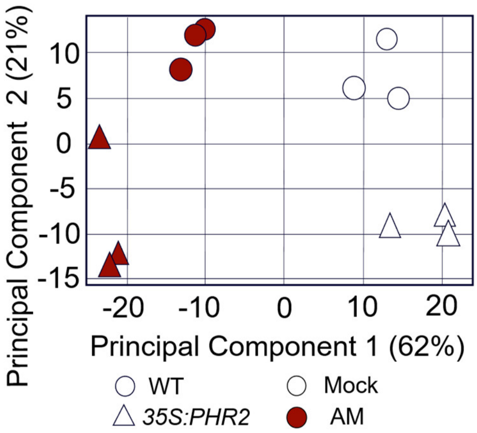

**Fig. S5.** PCA plot for the RNA-Seq based transcriptome of mock and AMF-inoculated wild-type and *35S:PHR2* roots grown at HP conditions.

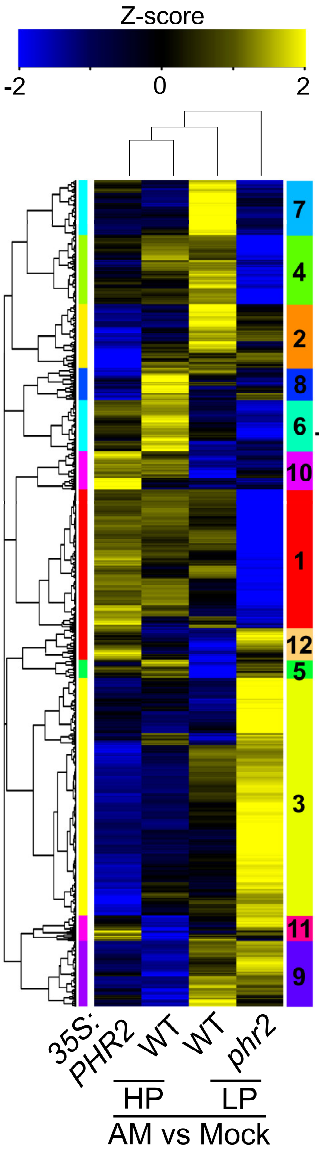

**Cluster 7 genes with positive Z-scores in *35S:PHR2* HP and WT LP samples but negative Z-scores in *phr2* LP and WT HP samples**

| Term | Description | neglogP |
|---|---|---|
| GO:0055114 | oxidation-reduction process | 4.44 |
| GO:0005975 | carbohydrate metabolic process | 3.94 |
| GO:0006629 | lipid metabolic process | 2.79 |
| GO:0006633 | fatty acid biosynthetic process | 2.47 |
| GO:0016114 | terpenoid biosynthetic process | 2.12 |
| GO:0006952 | defense response | 1.98 |
| GO:0008219 | cell death | 1.90 |
| GO:0009239 | enterobactin biosynthetic process | 1.86 |
| GO:0006915 | apoptotic process | 1.85 |
| GO:0008610 | lipid biosynthetic process | 1.83 |
| GO:0009607 | response to biotic stimulus | 1.82 |
| GO:0055085 | transmembrane transport | 1.68 |
| GO:0008299 | isoprenoid biosynthetic process | 1.65 |
| GO:0006631 | fatty acid metabolic process | 1.31 |
| KEGG:osa00900 | Terpenoid backbone biosynthesis | 1.29 |

**Cluster 1 DEGs with positive Z-scores in *35S:PHR2* HP and WT LP samples but negative Z-scores in *phr2* LP and WT HP samples**

| Term | Description | neglogP |
|---|---|---|
| GO:0016310 | phosphorylation | 5.35 |
| GO:0009875 | pollen-pistil interaction | 4.96 |
| GO:0006468 | protein phosphorylation | 4.14 |
| GO:0048544 | recognition of pollen | 3.56 |
| GO:0019748 | secondary metabolic process | 3.11 |
| GO:0006952 | defense response | 3.08 |
| GO:0055114 | oxidation-reduction process | 3.05 |
| GO:0055085 | transmembrane transport | 3.04 |
| GO:0009607 | response to biotic stimulus | 2.91 |
| GO:0006636 | unsaturated fatty acid biosynthetic process | 2.23 |
| GO:0006629 | lipid metabolic process | 2.16 |
| GO:0046686 | response to cadmium ion | 2.00 |
| GO:0019288 | isopentenyl diphosphate biosynthetic process, methylerythritol 4-phosphate pathway | 2.00 |
| GO:0006091 | generation of precursor metabolites and energy | 1.94 |
| GO:0010218 | response to far red light | 1.92 |
| GO:0006811 | ion transport | 1.88 |
| GO:0009744 | response to sucrose | 1.68 |
| GO:0006865 | amino acid transport | 1.68 |
| GO:0006813 | potassium ion transport | 1.62 |
| GO:0016117 | carotenoid biosynthetic process | 1.51 |
| GO:0019344 | cysteine biosynthetic process | 1.48 |
| GO:0080167 | response to karrikin | 1.47 |
| GO:0015995 | chlorophyll biosynthetic process | 1.45 |
| GO:0006520 | cellular amino acid metabolic process | 1.39 |
| GO:0003333 | amino acid transmembrane transport | 1.38 |
| GO:0006655 | phosphatidylglycerol biosynthetic process | 1.36 |
| KEGG:osa00904 | Diterpenoid biosynthesis | 2.23 |
| KEGG:osa00400 | Phenylalanine, tyrosine and tryptophan biosynthesis | 1.77 |

**Fig. S6.** Hierarchical clustering of combined DEGs (AM vs Mock samples, $\log_2$(Fold-change), lfc) from roots of *phr2* and *35S:PHR2* in LP and HP respectively and wild type at both $P_i$ conditions. Z-scores represent scaled lfc. Colored bars on the left side of heatmap depict individual clusters (based on the dendrogram). Gene ontology (GO) enrichment analysis for selected DEGs in cluster-1 and -7, which showed positive Z-scores for *35S:PHR2* HP and WT LP samples and negative Z-scores for *phr2* LP and WT HP samples. GO term enrichment indicates functional categories important for AM symbiosis such as fatty acid, lipid and carotenoid metabolism, response to biotic stimulus and response to karrikin.

A

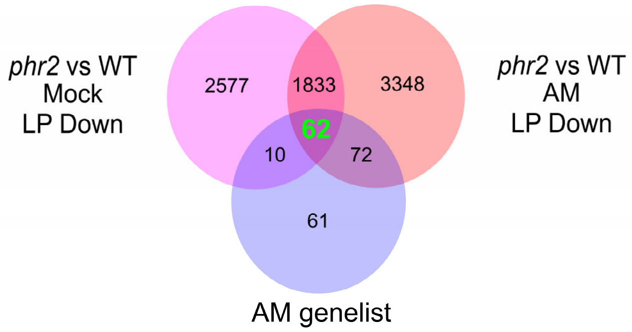

B

| Locus ID (62 Genes) | Gene description |
|---|---|
| **LOC_Os01g65000** | **AMT3;1, ammonium transporter protein** |
| **LOC_Os03g45290** | **Ankyrin repeat domain-containing protein (Medtr6g027840 VAPYRIN)** |
| **LOC_Os01g54270** | **D10, CCD8B** |
| **LOC_Os01g38580** | **D10-like, CCD8A** |
| **LOC_Os04g46470** | **D17, CCD7, carotenoid cleavage dioxygenase 7** |
| **LOC_Os11g01050** | **EXO70 exocyst complex subunit domain containing protein** |
| **LOC_Os03g29480** | **NSP1, GRAS transcription factor, nodulation-signaling pathway 1 protein** |
| **LOC_Os03g15680** | **NSP2, nodulation-signaling pathway 2 protein** |
| **LOC_Os05g41090** | **OsDMI3, OsCCaMK, calcium/calmodulin depedent protein kinases** |
| **LOC_Os06g02520** | **OsIPD3, OsCYCLOPS, interacting protein of DMI3** |
| **LOC_Os03g13080** | **OsLysM-RLK2,OsLYK5, MYR1/LYK2/RLK2/NFR5** |
| **LOC_Os06g44430** | **Protein kinase (Medtr4g129010 KIN2)** |
| **LOC_Os01g46860** | **PT11, inorganic phosphate transporter** |
| **LOC_Os04g10800** | **PT13, inorganic phosphate transporter** |
| **LOC_Os09g23640** | **STR1, ABC-2 type transporter domain containing protein** |
| **LOC_Os07g38070** | **SYMRK, protein kinase, putative, expressed (Lj2g3v1467920 LjSymRK)** |
| **LOC_Os09g15240** | **Zaxinone Synthase (ZAS), carotenoid cleavage dioxygenase** |
| LOC_Os08g05690 | ABC transporter, ATP-binding protein, putative |
| LOC_Os05g50300 | AMP1, AMP-binding enzyme, putative |
| LOC_Os04g39780 | AMP-binding enzyme family protein |
| LOC_Os01g44950 | AMP-binding enzyme, putative |
| LOC_Os07g38750 | AP2 domain containing protein |
| LOC_Os12g39330 | AP2 domain containing protein |
| LOC_Os02g48820 | BCP, plastocyanin-like domain containing protein, putative, expressed |
| LOC_Os06g47130 | C2domain containing protein |
| LOC_Os07g23450 | C2H2 zinc finger protein |
| LOC_Os08g28240 | carotenoid cleavage dioxygenase, putative |
| LOC_Os04g58680 | CBF1/2, core histone H2A,H2B,H3,H4, putative |
| LOC_Os06g20120 | CND41, chloroplast nucleoid DNA binding protein, putative, expressed |
| LOC_Os09g39530 | Cupin-domain contaning protein |
| LOC_Os07g33620 | cytochrome P450 domain containing protein |
| LOC_Os01g50520 | cytochrome P450 monooxygenase CYP711A12, putative, expressed |
| LOC_Os01g50590 | cytochrome P450, putative, expressed |
| LOC_Os05g51240 | D14L2a, Hydrolase, alpha/beta fold family domain containing protein |
| LOC_Os10g42400 | DNA polymerase III, clamp loader complex, gamma/delta/delta subunit |
| LOC_Os01g46390 | DUF538 domain containing protein, putative |
| LOC_Os05g49790 | DUF538 domain containing protein, putative |
| LOC_Os08g42990 | expressed protein |
| LOC_Os11g29630 | expressed protein |
| LOC_Os02g18954 | GDSL-like lipase,acylhydrolase, putative |
| LOC_Os02g44850 | GDSL-like lipase,acylhydrolase, putative |
| LOC_Os04g47390 | GDSL-like lipase,acylhydrolase, putative |
| LOC_Os07g19040 | glycosyl hydrolase, putative |
| LOC_Os08g34258 | inhibitor I family protein, putative |
| LOC_Os08g34249 | inhibitor I family protein, putative, expressed |
| LOC_Os04g39180 | KIN6, nodulation receptor kinase precursor, putative |
| LOC_Os02g20140 | KinD, protein kinase domain containing protein |
| LOC_Os01g57400 | lysM domain containing protein, putative |
| LOC_Os04g40570 | MDR1, ABC transporter, ATP-binding protein, putative |
| LOC_Os05g47500 | MDR-like ABC transporter |
| LOC_Os08g42590 | mtN19, putative, expressed |
| LOC_Os01g52750 | OsSub3 - Putative Subtilisin homologue |
| LOC_Os04g04750 | peroxidase precursor, putative, expressed |
| LOC_Os03g57740 | plastocyanin-like domain containing protein, putative |
| LOC_Os02g57700 | protein kinase, putative, expressed |
| LOC_Os01g72710 | putative RETICULATA-RELATED protein of unknown function |
| LOC_Os09g20970 | receptor kinase, putative |
| LOC_Os03g38600 | secretory carrier-associated membrane protein, putative, expressed |
| LOC_Os08g29510 | Taurine catabolism dioxygenase TauD/TfdA domain containing protein |
| LOC_Os10g18510 | UDP-glucoronosyl and UDP-glucosyl transferase domain containing protein |
| LOC_Os05g50680 | WRKY83 |
| LOC_Os01g53130 | zinc finger, C3HC4 type domain containing protein, expressed |

**Fig. S7. Overlap of genes with decreased transcript levels in *phr2* in Mock or AM roots with AM genelist.** (**A**) Venn diagram of DEGs downregulated in *phr2* vs WT Mock and AM roots at LP and AM genelist. (**B**) Genes common to all the three sets. Genes highlighted in red have been previously genetically shown to be required for AM development or function.

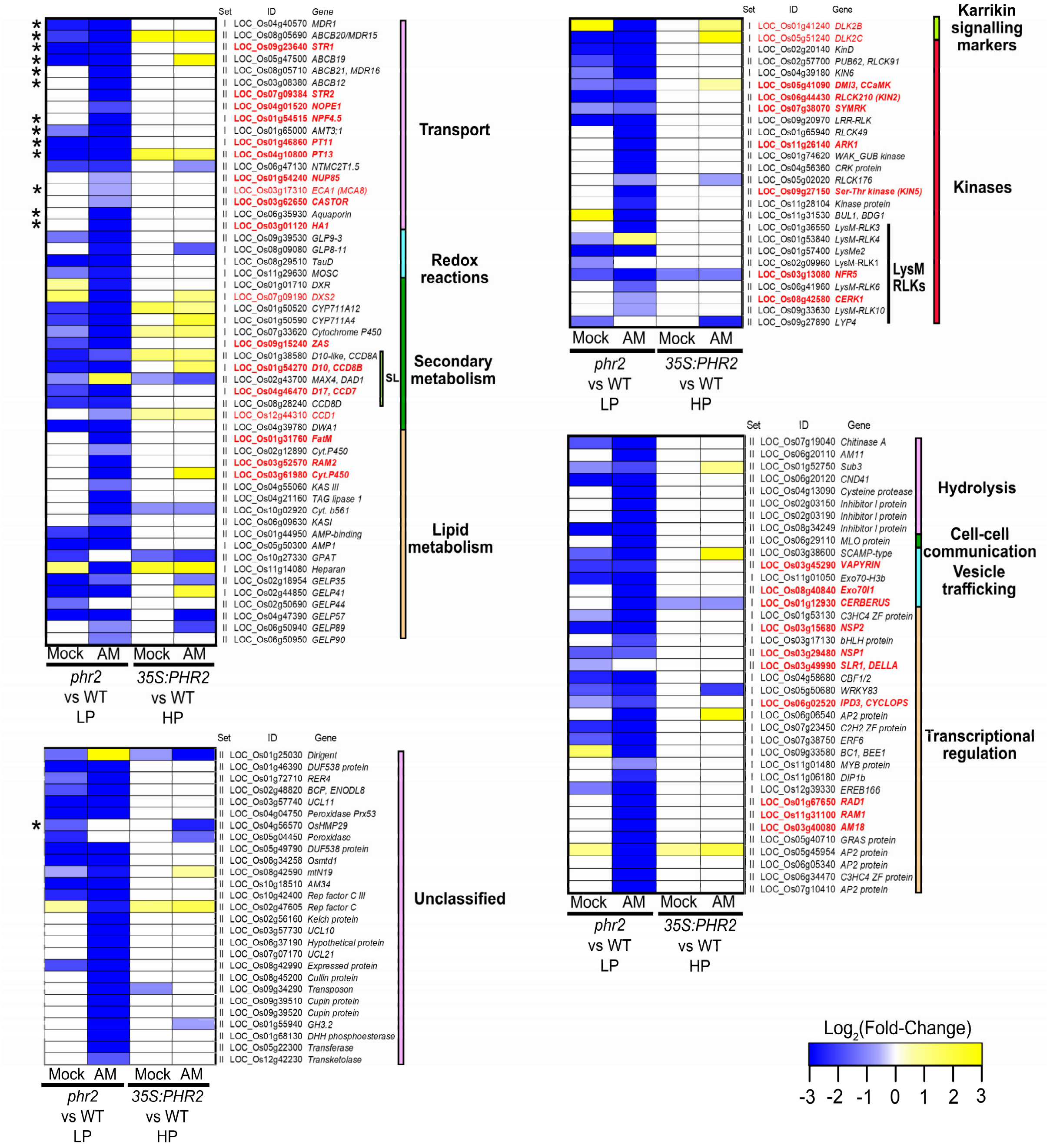

**Fig. S8. Expression of genes required for or induced during AM depend on PHR2.** Heatmaps for $\log_2$(Fold-change) of AM genes from set I and II (Figure 2G) for the comparisons *phr2* vs wild type at LP (25 µM) and *35S:PHR2* vs wild type at HP (500 µM). Colored bars on the right indicate functional categories to which the genes belong. Genes with genetically confirmed functions in AM symbiosis are indicated in red (bold for mutants, regular font for *RNAi* lines). 144 out of 205 genes (70%) in the AM genelist had reduced expression in *phr2* vs WT AM and/or Mock LP samples. Asterisks indicate orthologs of *Lotus japonicus* genes involved in transport and associated with P1BS elements in Supplementary table 2 of Lota et al., 2013. Out of 48 total genes in this table, only 31 could be retrieved from the *Lotus japonicus* genome assembly build 1.0 (http://www.kazusa.or.jp/lotus/release1/) and 14 out of these have reduced expression in rice *phr2* vs WT).

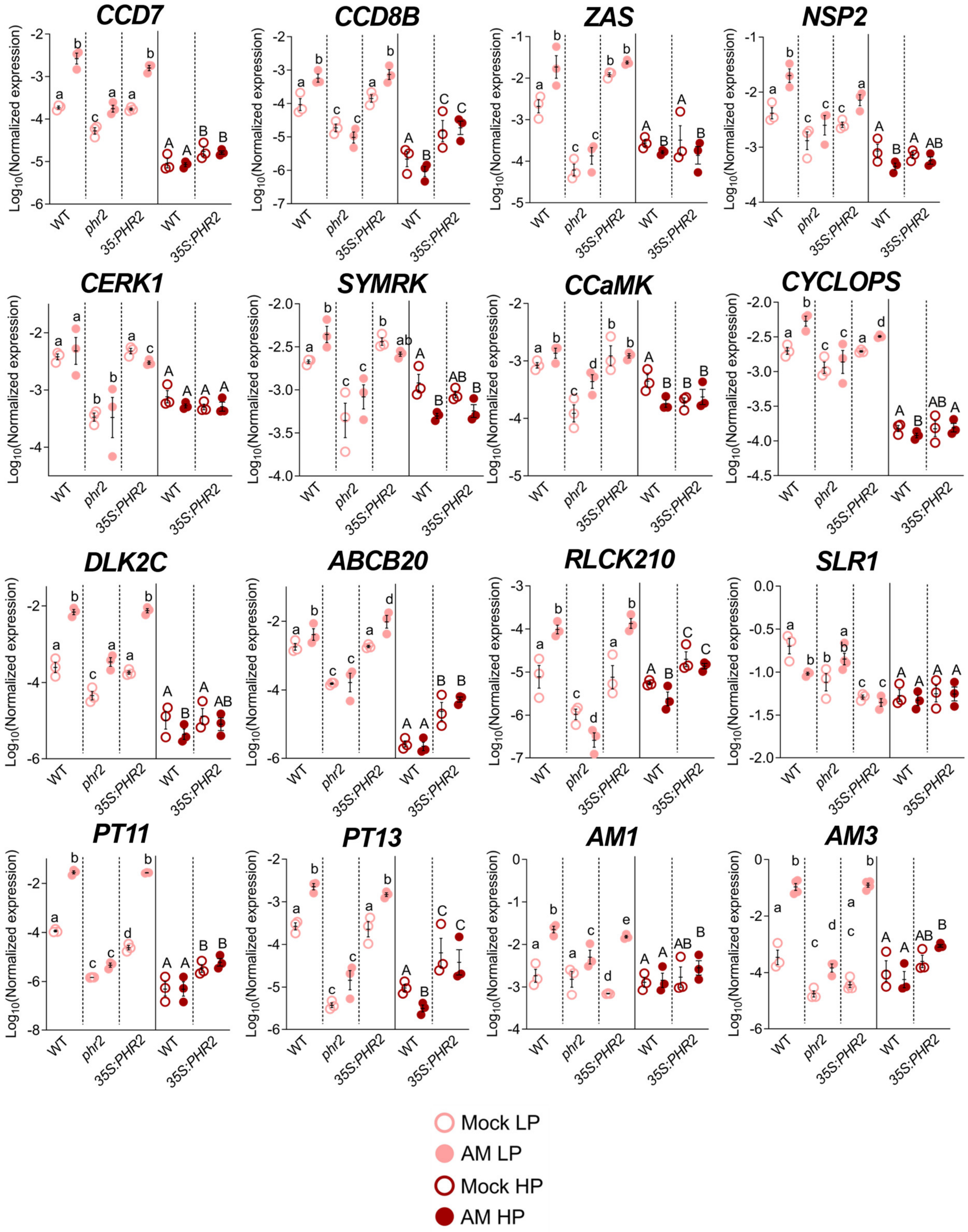

**Fig. S9. RT-qPCR-based transcript accumulation of selected DEGs recapitulates the RNA-Seq results.** Relative transcript accumulation in mock inoculated (Mock) and *R. irregularis* colonized (AM) roots of the indicated genotypes grown in quartz sand and fertilized with LP (25 µM $P_i$) or HP (500 µM $P_i$). Expression values of indicated genes were normalized to the geometric mean of the expression of two housekeeping genes, *UBIQUITIN* and *CYCLOPHILIN*. Letters indicate statistical differences between genotypes and treatments for each phosphate level separately. Statistics: Individual data-points and mean ± SE are shown. N=3. Brown-forsythe and Welch’s One-Way ANOVA test with Games-Howell's multiple comparisons test was carried out for datapoints between samples at each phosphate level separately. Different letters indicate statistical differences between samples.

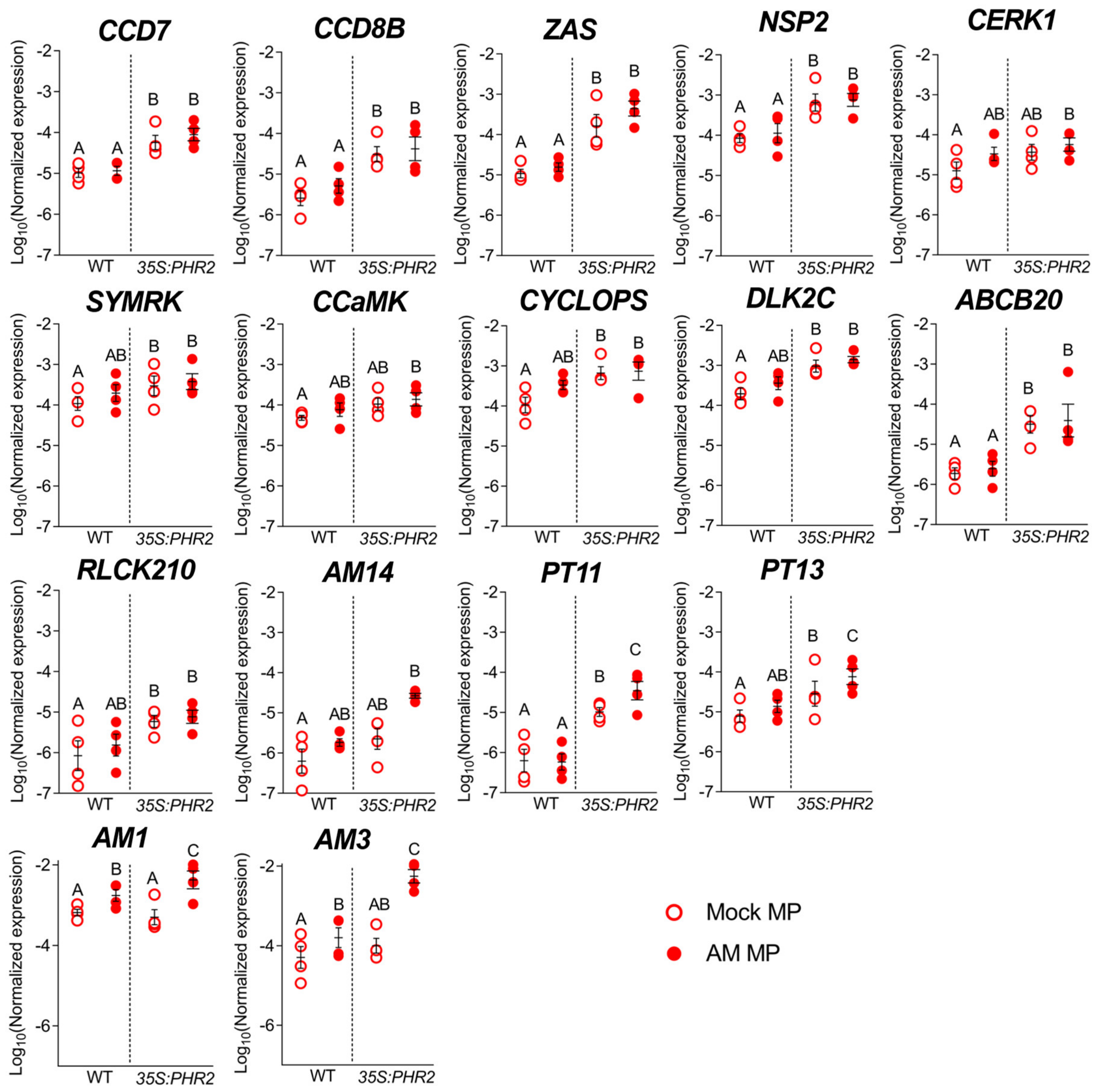

**Fig. S10. RT-qPCR-based transcript accumulation of selected DEGs at medium phosphate.** Relative transcript accumulation in mock inoculated (Mock) and *R. irregularis* colonized (AM) roots of the indicated genotypes grown in the same experiment as Fig. S3 in quartz sand and fertilized with MP is shown. Expression values of indicated genes were normalized to the geometric mean of the expression of two housekeeping genes, *UBIQUITIN* and *CYCLOPHILIN*. Letters indicate statistical differences between genotypes and treatments within each phosphate level. Statistics: Individual data-points and mean ± SE are shown. N=3-4. Brown-forsythe and Welch's One-Way ANOVA test with Games-Howell's multiple comparisons test was carried out for datapoints between all samples. Different letters indicate statistical differences between samples.

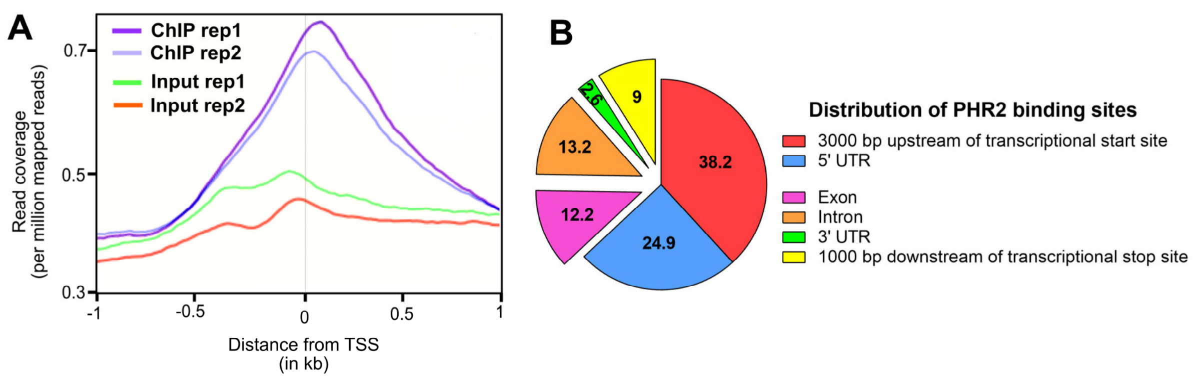

**Fig. S11. ChIP-Seq binding peaks of PHR2-FLAG are enriched near the transcriptional start site**. (**A**) Read coverage plot for the two biological replicates from ChIP-Seq with FLAG tagged PHR2 protein. TSS, transcriptional start site. (**B**) Distribution of PHR2-binding sites in the rice genome. ChIP-Seq read distribution in relation to transcriptional start site (TSS) suggested a slight skew in the distribution of PHR2 binding sites towards 1000 bp downstream of TSS. Correspondingly, PHR2 binding sites are enriched not only in 3000 bp region upstream of TSS (38.2%) but also in the regions downstream of TSS such as 5′ UTR, exon and intron (24.9% + 12.2% + 13.2% = 50.3%).

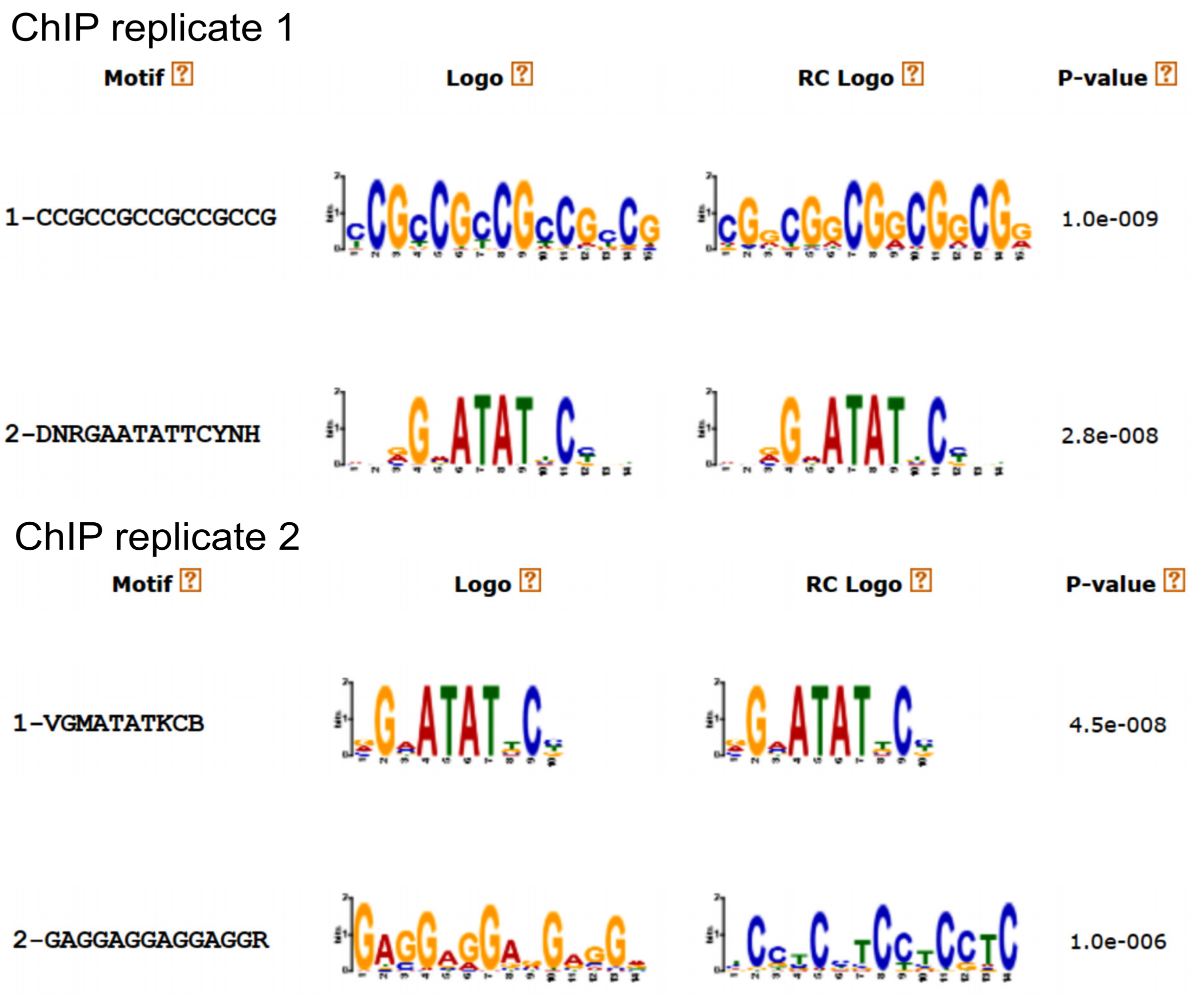

**Fig. S12.** Motifs over-represented in DNA sequences with PHR2 binding sites. Analysis was carried out separately for the two biological ChIP-Seq replicates using STREME (https://meme-suite.org/meme/tools/streme).

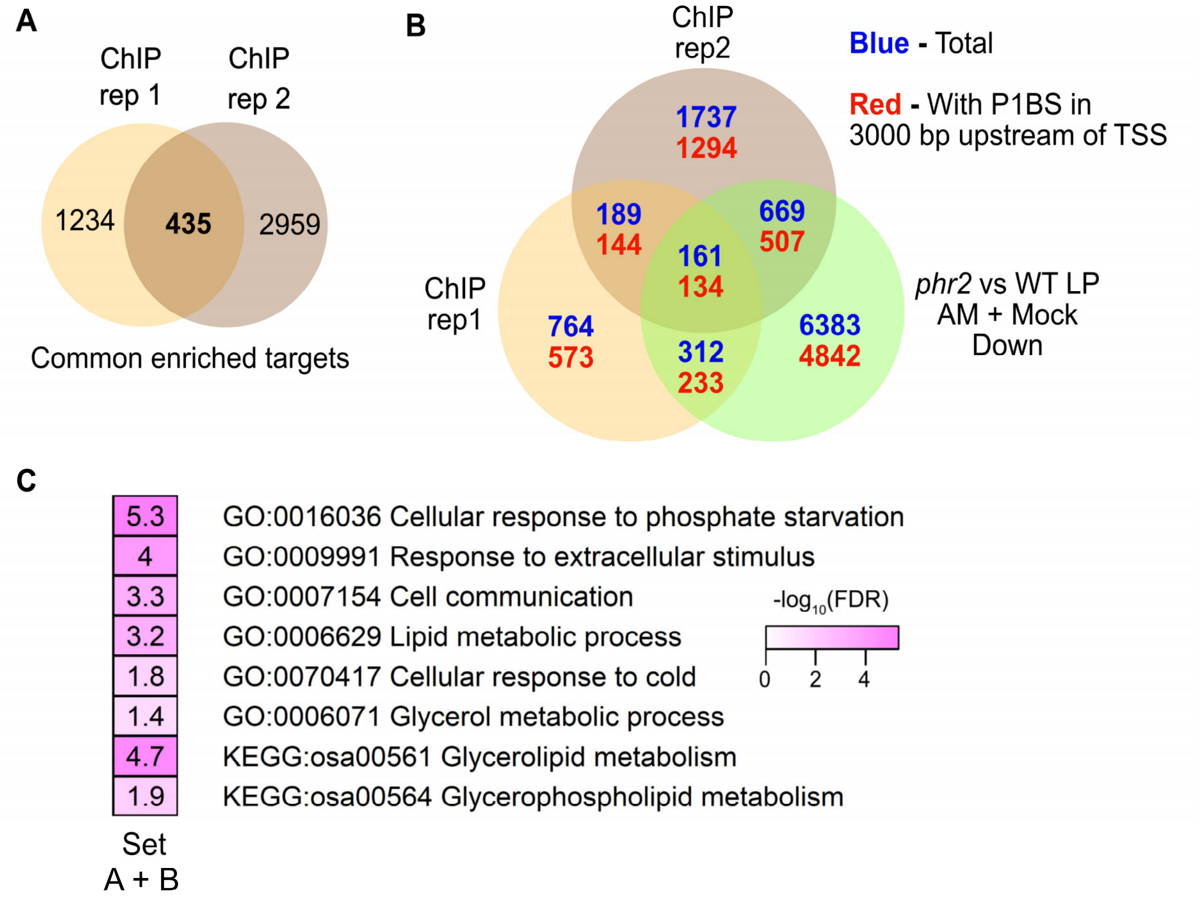

**D**

| Enriched motifs (Set A + B) | E-value | Significance |
|---|---|---|
| GAATATGC \| GCATATTC | 1.00E-05 | 4.99 |
| GAATATCC \| GGATATTC | 5.00E-05 | 4.3 |

**Fig. S13. Binding site analysis for rice PHR2.** (**A**) Venn diagram showing overlap of PHR2 targets from two independent biological ChIP-Seq replicates. These 435 common PHR2 targets are genes annotated closest to PHR2-binding sites in both replicates. (**B**) Venn diagram showing overlap of PHR2 targets with DEGs with reduced expression in *phr2* vs WT AM + Mock samples at LP. Blue indicates the total number of genes and red with P1BS or P1BS-like motif in 3000 bp upstream region upstream of transcriptional start site (TSS)). MSU IDs in the three individual gene sets were converted to RAPDB Locus IDs (to facilitate extraction of upstream sequence from RAPDB website, https://rapdb.dna.affrc.go.jp/). This resulted in a smaller number of genes than the original number of MSU ID DEGs. (**C**) GO-term enrichment in category "biological process" for Set A + Set B genes (167 genes out of 435 PHR2 targets which are repressed in AM or Mock root samples of *phr2* vs WT grown at LP as shown in Fig. 3A). Darker colors indicate stronger enrichment of GO-terms. GO-terms include categories involved in phosphate starvation signalling as well as AM. (**D**) Motifs enriched in 1000 bp sequence (upstream of TSS) for Set A + B genes in Fig. 3A.

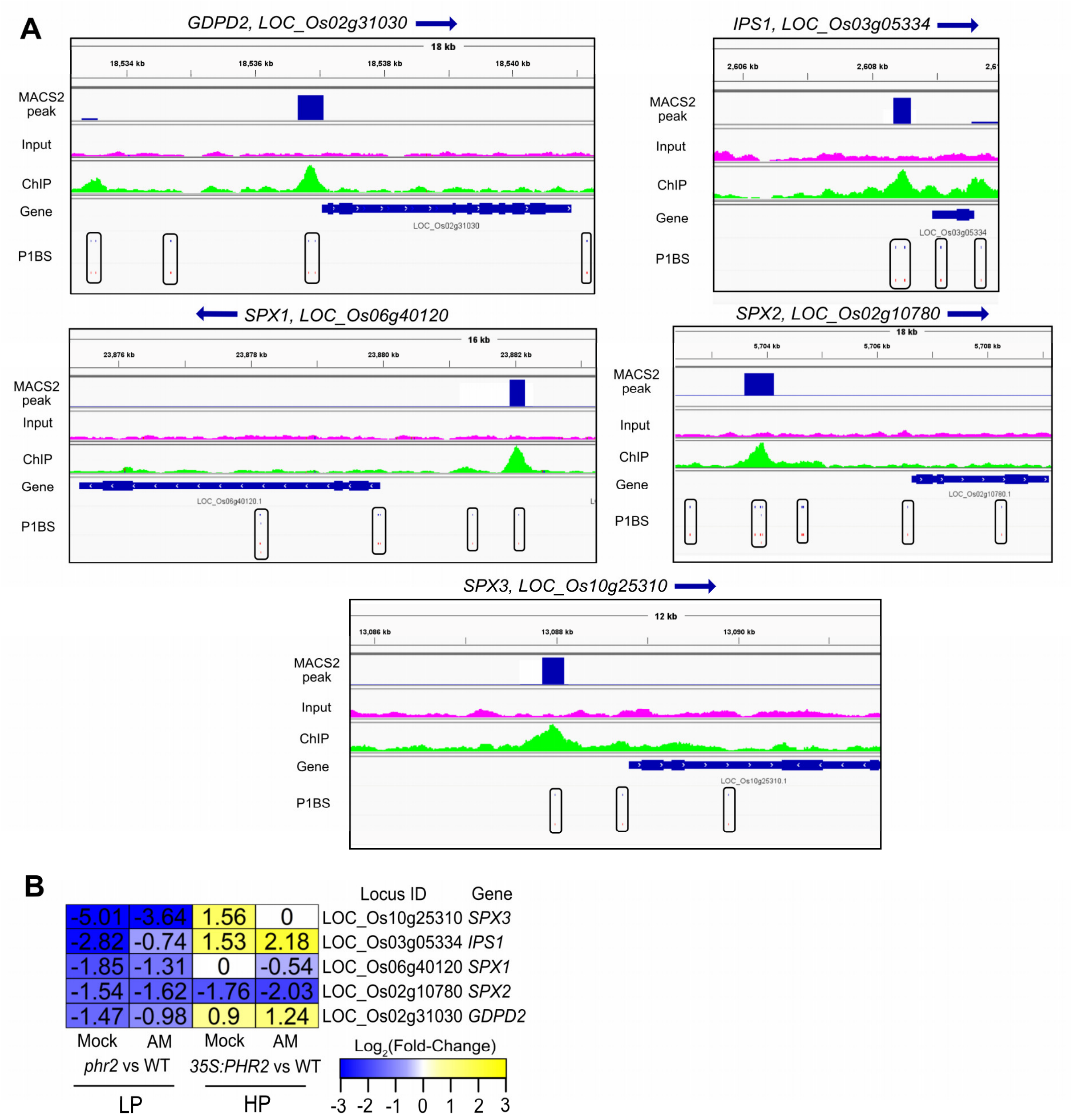

**Fig. S14. IGV browser view of ChIP-Seq peaks adjacent to previously known PHR2 target genes.** (**A**) ChIP-Seq peak profiles of genes which have been previously shown to be PHR2 targets. Gene orientation is indicated by the direction of the blue arrow close to the gene name. MACS2 peaks (blue bars) denote PHR2 binding sites corresponding to enrichment of PHR2-FLAG IP (green color) sequencing reads vs Input (pink color) sequencing reads. Positions of P1BS elements along the genomic coordinate are marked by enclosing the motifs in black rectangular boxes. (**B**) RNA-Seq based $log_2$(Fold-change) of these known PHR2 target genes in for *phr2* vs wild type and *35S:PHR2* vs wild type at LP (25 μM) and HP (500 μM), respectively. The phosphate level at HP (500 μM) maybe high enough to prevent the transcriptional induction of some of these phosphate starvation response genes such as *SPX1* and *2* in the *35S:PHR2* line.

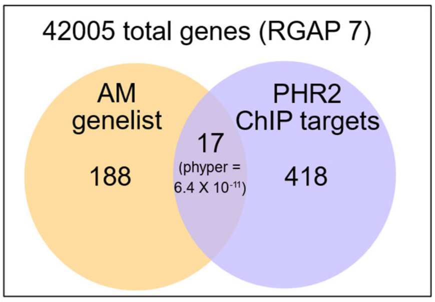

**Fig. S15. Enrichment of PHR2 direct targets in AM genelist.** Hypergeometric test was used to assess the statistical significance (phyper) of overlap of PHR2 ChIP targets with the AM genelist.

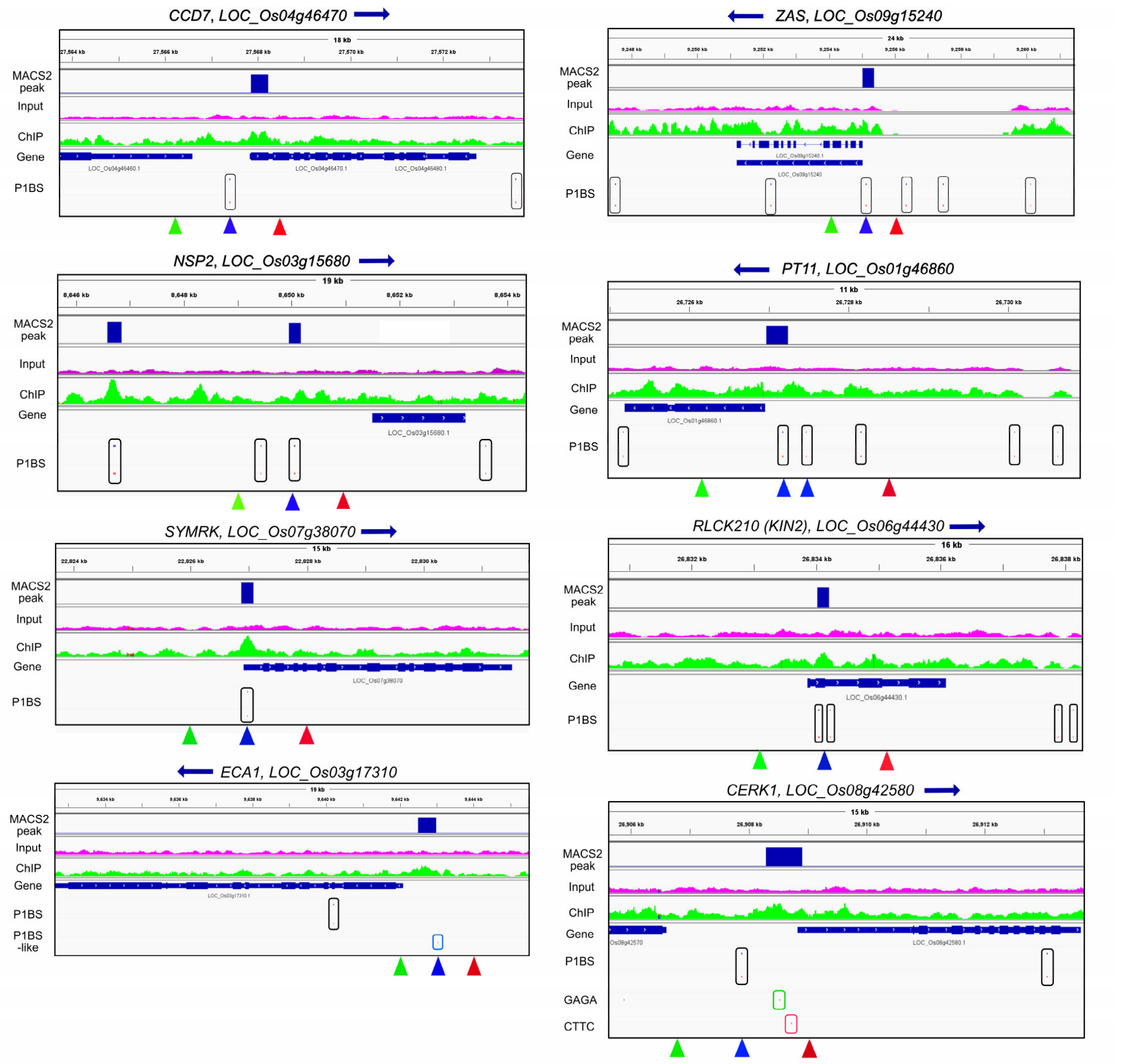

**Fig. S16. IGV browser view of ChIP-Seq peaks adjacent to AM-relevant genes.** The selected genes are those depicted in red in Fig. 3C. Gene orientation is indicated by the direction of the blue arrow next to the gene name. MACS2 peaks (blue bars) indicate PHR2 binding sites corresponding to enrichment of PHR2-FLAG IP (green color) sequencing reads vs Input (pink color) sequencing reads. The positions of P1BS elements along the genomic coordinate are marked by enclosing the motifs in black rectangular boxes. In the *ECA1* promoter, the P1BS-like motif is marked by a blue rectangular box. In the *CERK1* promoter, the GAGA and CTTC motifs are marked by enclosing them in green and red rectangular boxes, respectively. Blue triangles indicate the position of ChIP-qPCR (Fig. S15) primers flanking the P1BS motifs, while green and red triangles indicate the position of primers 1000 bp left (5’) of P1BS (-1000 bp) and +1000 bp right (3’) of P1BS (+1000 bp), respectively. Motifs: P1BS is GNATATNC; P1BS-like is AMATATYC; GAGA is GGAGAGGA; CTTC is TCCTCTTGTTCTTC.

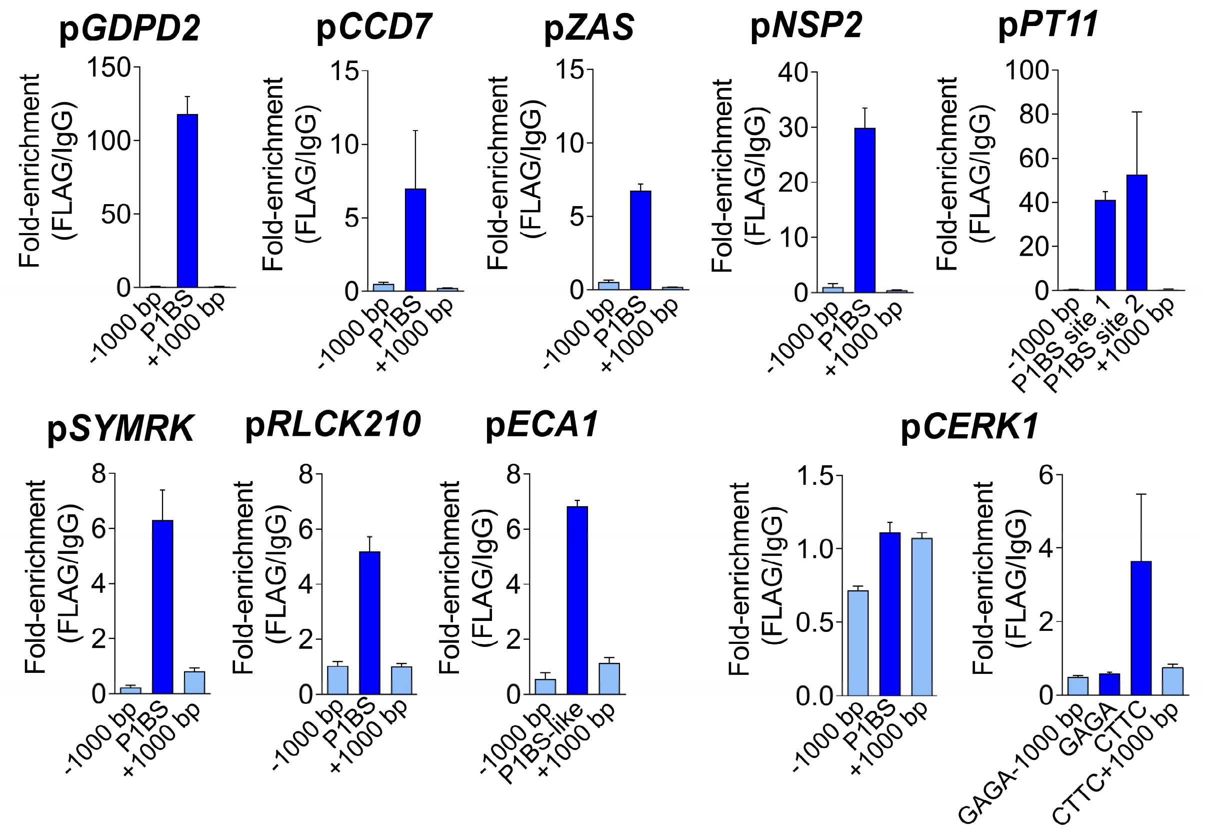

**Fig. S17. Enrichment of PHR2 at P1BS promoter motifs detected by ChIP-qPCR.** Primers were designed to amplify regions flanking motifs (P1BS, P1BS-like, GAGA, CTTC), and 1000 bp left (5') of these motifs (-1000 bp) and 1000 bp right (3') of these motifs (+1000 bp). Motifs: P1BS is GNATATNC; P1BS-like is AMATATYC; GAGA is GGAGAGGA; CTTC is TCCTCTTGTTCTTC. Data: Individual data-points and mean ± SE are shown. N=3.

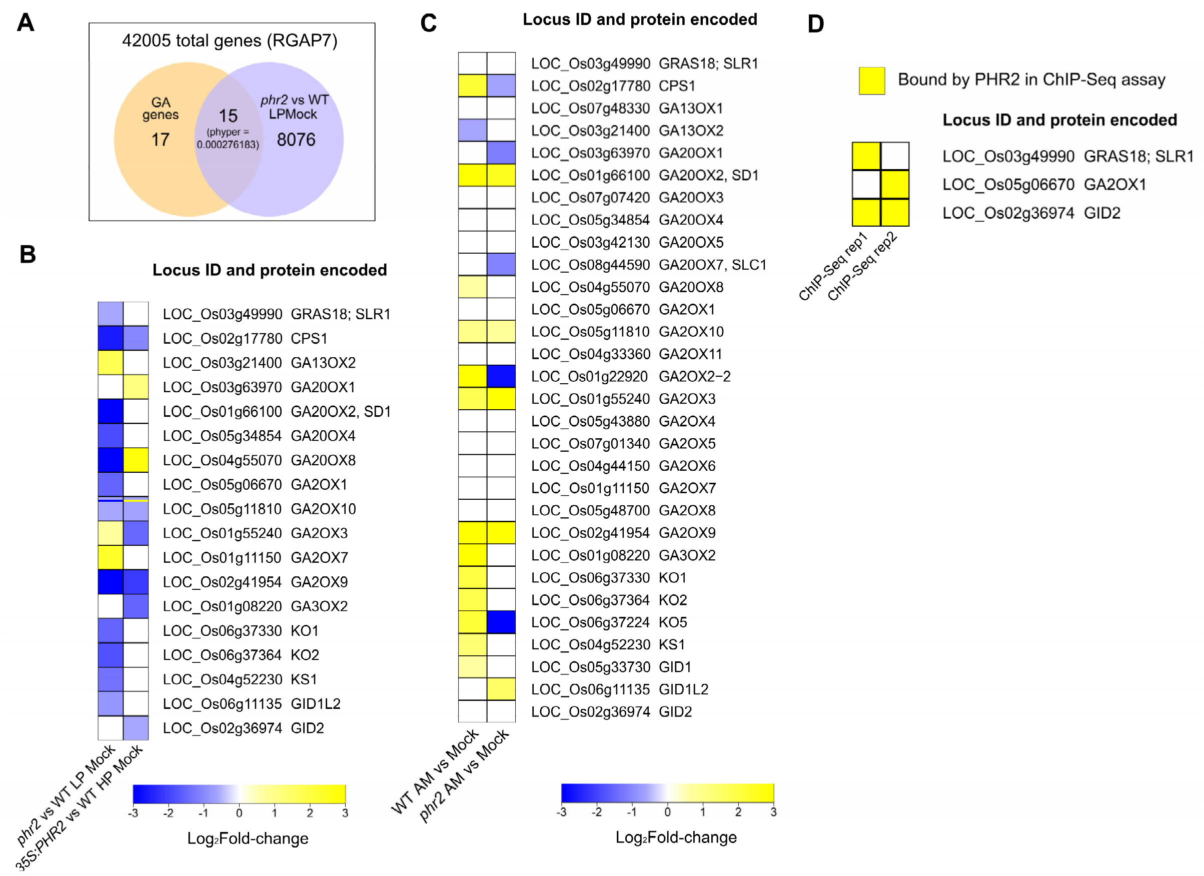

**Fig. S18. Gibberellin-biosynthesis and -signaling related genes in RNASeq and ChIP-Seq**. (**A**) Gibberellin (GA)-related genes are enriched in DEGs with reduced expression in non-inoculated *phr2* vs wild type. as shown by a hypergeometric test to assess the statistical significance (phyper). (**B**) Expression of GA-related genes in non-colonized *phr2* vs wild type roots at LP and *35S:PHR2* vs wild type roots at HP. (**C**) Comparison of gene expression for GA-related genes in *phr2* and wild type in AM vs Mock roots. (**D**) GA-related genes directly targeted by PHR2 as determined by ChIP-Seq.

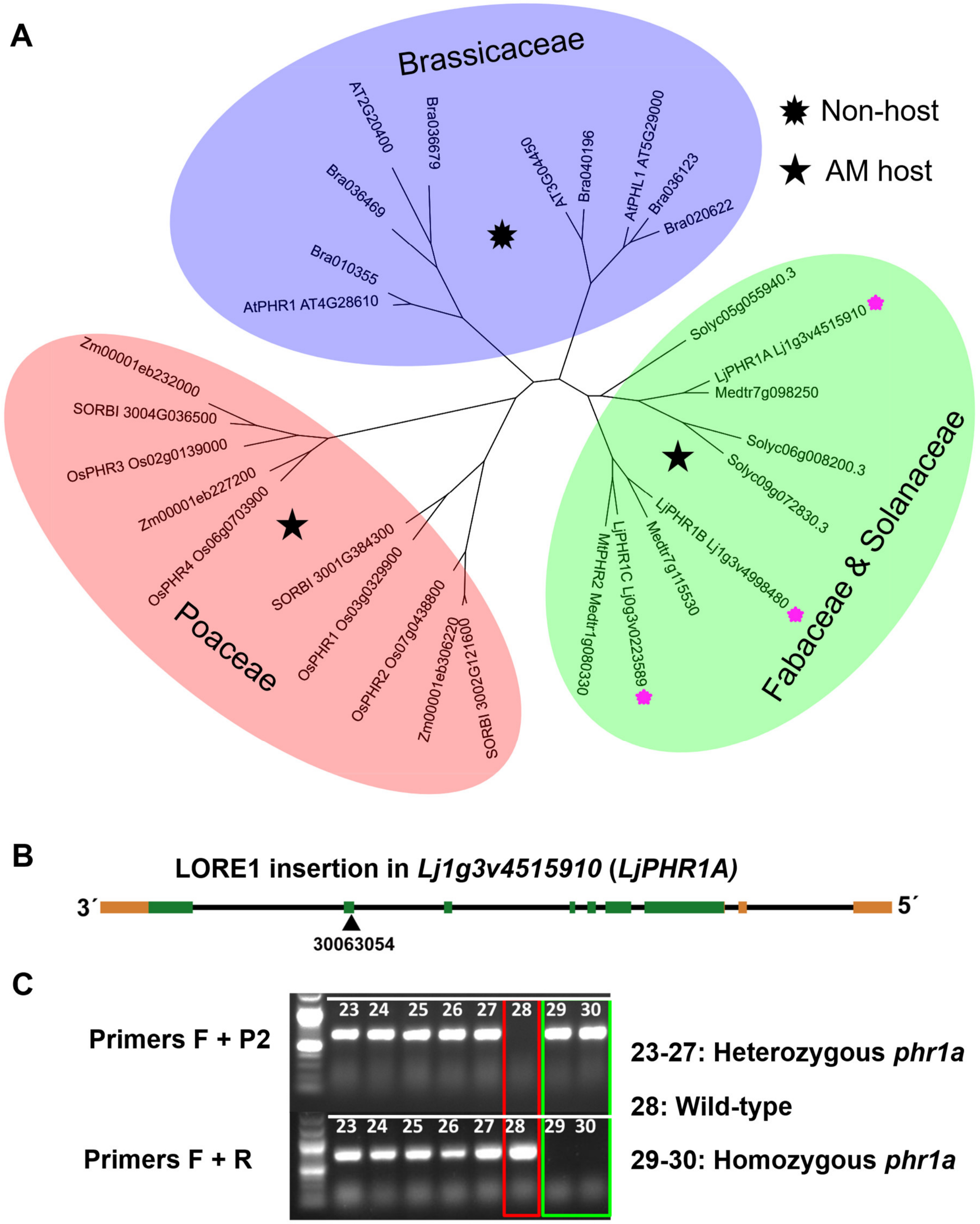

**Fig. S19. *Lotus japonicus* PHR1A protein.** (**A**) Phylogenetic tree of PHR proteins in representative *Brassicaceae, Poaceae, Fabaceae* and *Solanaceae*. The three *Lotus japonicus* PHR proteins are marked with pink stars. (**B**) Position of LORE1 insertion in *L. japonicus PHR1A*. The number indicates the Plant ID for LORE1 insertion. (**C**) Genotyping for LORE1 insertion in *phr1a*. The P2 primer sequence in located in the LORE1 insertion while F and R are *PHR1A* specific primers surrounding the insertion.

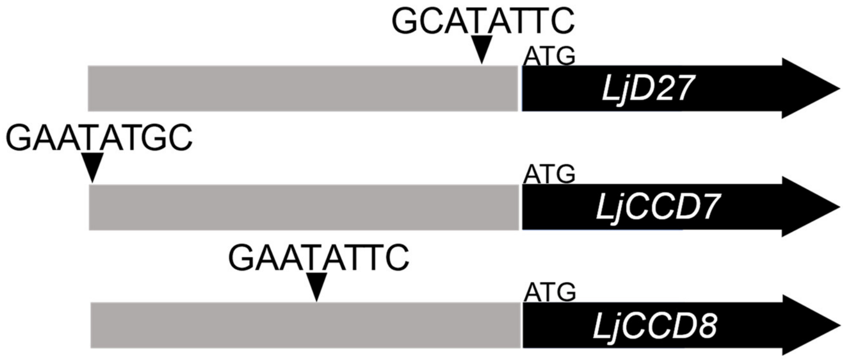

**Fig. S20. Position of P1BS motifs in the promoters of strigolactone biosynthesis genes in *Lotus japonicus***. Promoter of length 1600 kb is represented in gray for each gene.

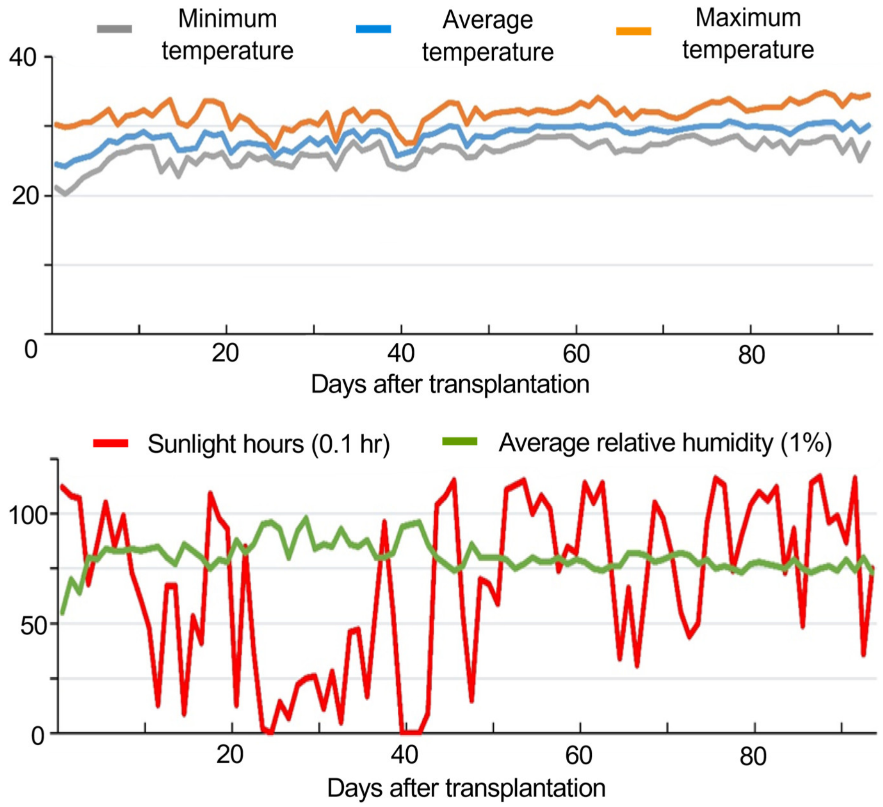

**Fig. S21.** Temperature, sunlight and relative humidity profiles during the greenhouse experiment in field soil.

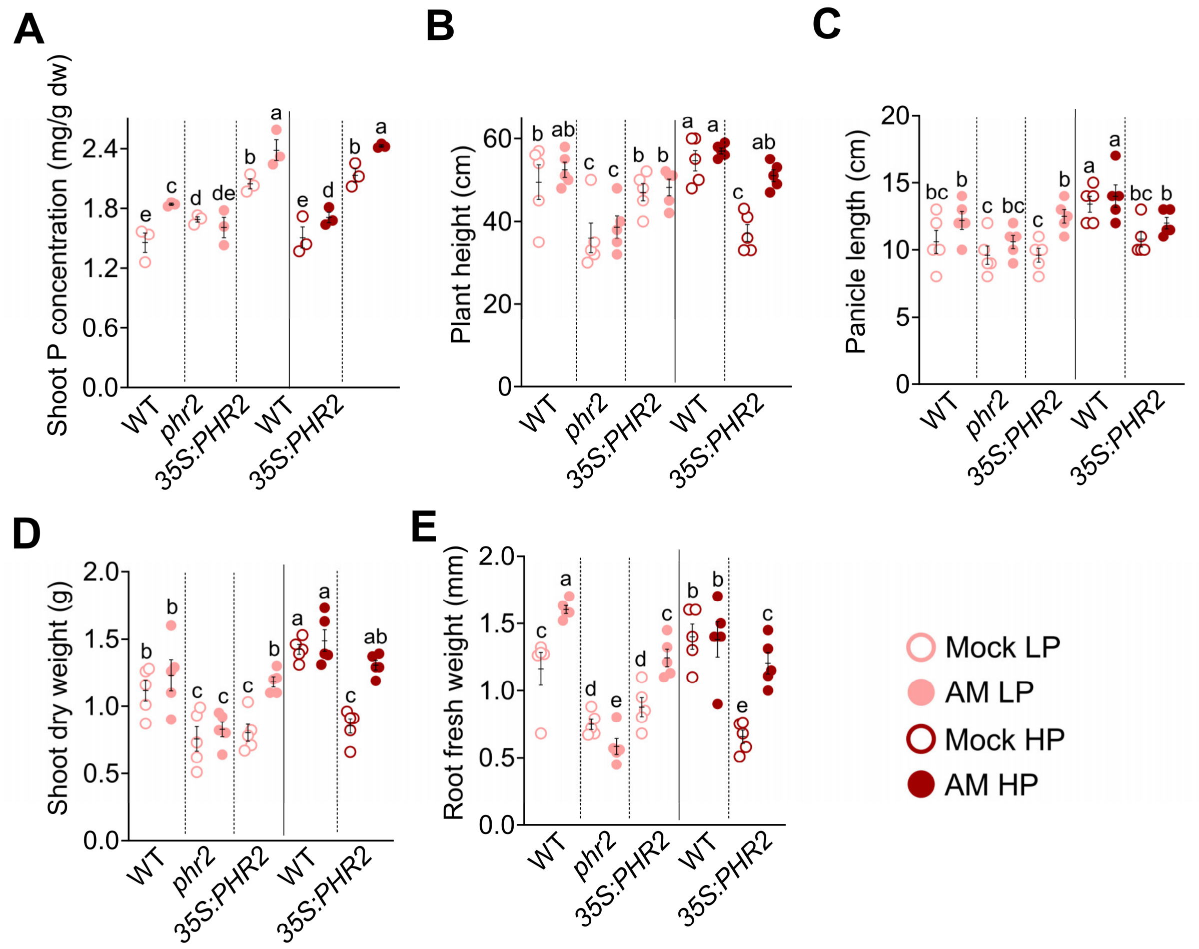

**Fig. S22. PHR2 affects rice agronomic traits in a field soil.** (**A**) Shoot phosphorus (P) concentration (mg/g dw), (**B**) Plant height (cm), (**C**) panicle length (cm), (**D**) shoot dry weight (g), (**E**) root fresh weight (g) in mock (Mock) and *R. irregularis* (AM) inoculated plants of wild type, *phr2* and *35S:PHR2* lines grown at LP (unfertilized) and of wild type and *35S:PHR2* lines grown at HP (fertilized with superphosphate fertilizer, $P_2O_5$). Traits were quantified for plants harvested at 110 days post transplanting into soil and inoculation. Statistics: Individual data-points and mean ± SE are shown. N=3-5; Brown-forsythe and Welch's One-Way ANOVA test with Games-Howell's multiple comparison test was carried out for datapoints between all samples. Different letters indicate statistical differences between samples.

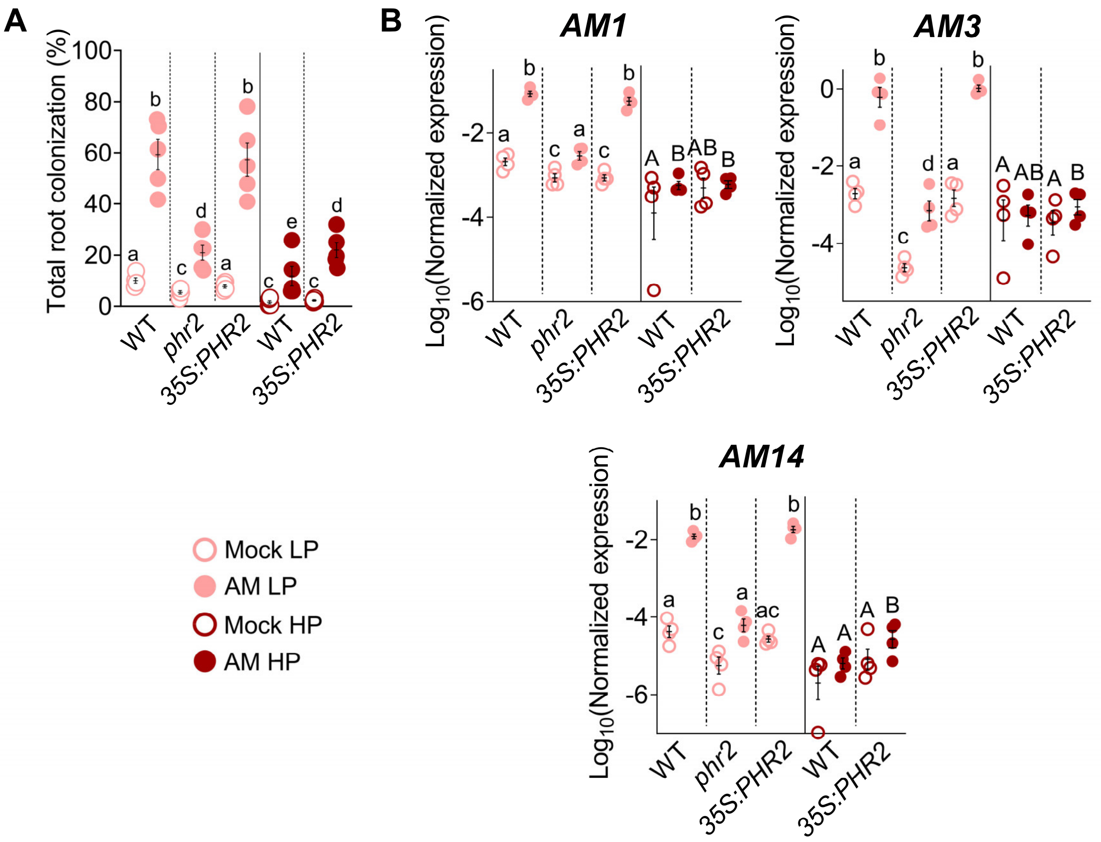

**Fig. S23. Root colonization and RT-qPCR-based transcript accumulation of AM-marker genes in roots of plants grown in field soil.** (**A**) Total root colonization (%) in *R. irregularis*-inoculated plants of wild type and *phr2* and *35S:PHR2* at LP (unfertilized) and in wild type and *35S:PHR2* lines at HP (fertilized with superphosphate fertilizer, $P_2O_5$). Roots were harvested at 110 days post transplantation into field soil and inoculation. Letters indicate statistical differences between genotypes, treatment and phosphate levels. Statistics: N=5; Kruskal-Wallis test with Dunn's posthoc comparison. (**B**) Relative transcript accumulation in mock inoculated (Mock) and *R. irregularis* colonized (AM) roots of the indicated genotypes grown in field soil and fertilized with LP or HP (as described in A). Expression values of indicated genes were normalized to the geometric mean of the expression of two housekeeping genes, *UBIQUITIN* and *CYCLOPHILIN*. Letters indicate statistical differences between genotypes and treatments within each phosphate level. Statistics: Individual data-points and mean ± SE are shown. (A) N=5; Brown-forsythe and Welch's One-Way ANOVA test with Games-Howell's multiple comparisons test between all samples. Different letters indicate statistical differences between samples. (B) N=3-4. Brown-forsythe and Welch's One-Way ANOVA test with Games-Howell's multiple comparisons test was carried out for datapoints between samples at each phosphate level separately. Different letters indicate statistical differences between samples.

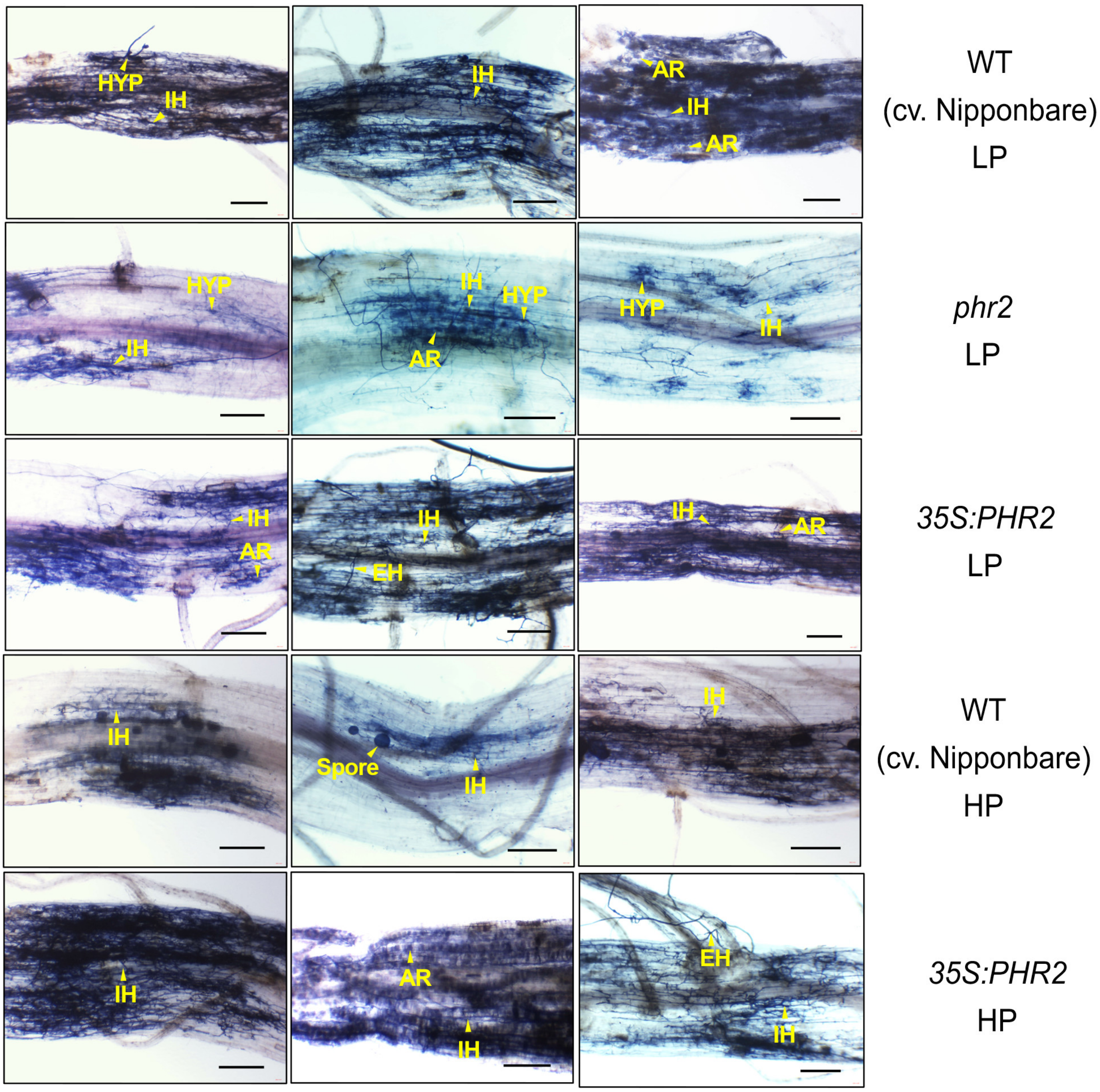

**Fig. S24.** Brightfield images of roots stained with acid-ink to visualize colonization of wild type (cv. Nipponbare), *phr2* and *35S:PHR2* roots by *R. irregularis* at 110 days post transplantation and grown at LP (unfertilized) or HP (fertilized with superphosphate fertilizer, $P_2O_5$) in field soil. Scale bars, 200 µm. Abbreviations: EH, extraradical hypha; HYP: hyphopodium; IH, intraradical hypha; AR, arbuscule, VE, vesicle.

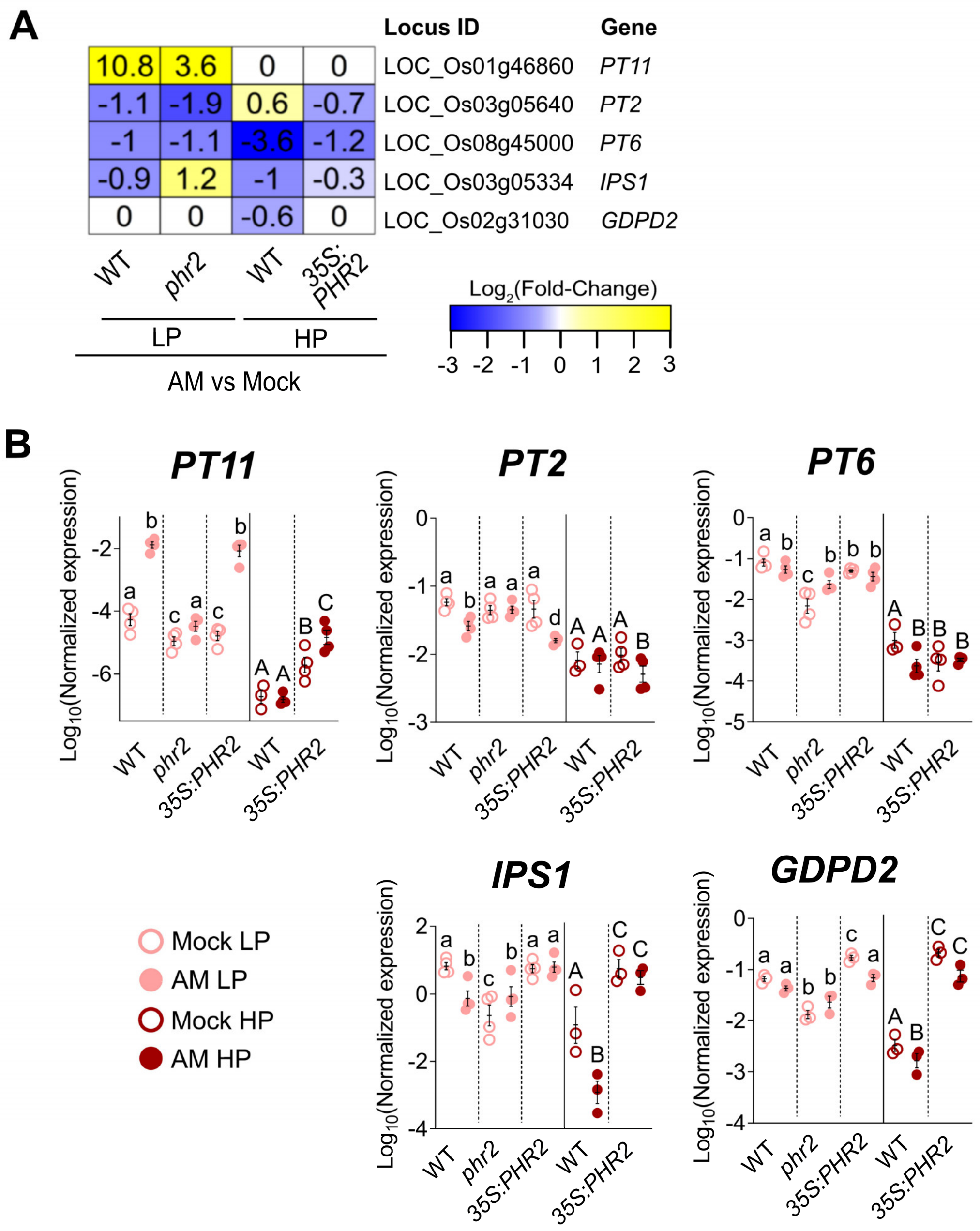

**Fig. S25. RT-qPCR-based transcript accumulation of phosphate transporters and starvation marker genes in roots of plants grown in field soil.** (**A**) RNA-seq-based fold-change of transcript accumulation (AM vs mock) of phosphate transporter genes involved in AM-mediated (*PT11*) or direct (*PT2*, *PT6*) $P_i$ uptake, as well as the phosphate starvation marker genes (*IPS1*, *GDPD2*) at LP (25 µM $P_i$) or HP (500 µm $P_i$) in quartz sand. (**B**) RT-qPCR based relative transcript accumulation in mock inoculated (Mock) and *R. irregularis* colonized (AM) roots of the indicated genotypes grown in field soil and fertilized with LP or HP (as described in A) is shown. Roots were harvested at 110 days post transplanting and inoculation. Expression values of indicated genes were normalized to the geometric mean of the expression of two housekeeping genes, *UBIQUITIN* and *CYCLOPHILIN*. Letters indicate statistical differences between genotypes and treatments within each phosphate level. Statistics: Individual data-points and mean ± SE are shown. N=3-4. Brown-forsythe and Welch's One-Way ANOVA test with Games-Howell's multiple comparisons test was carried out for datapoints between samples at each phosphate level separately. Different letters indicate statistical differences between samples.

**Table S1. Primers used for RT-qPCR and genotyping.**

| Primer name | Forward Primer (5'→3') | Reverse Primer (5'→3') |
|---|---|---|
| CCD7-qRT | AATGCACTTGTGGCAAAACTAGAG | CATTGGAAAAGTGAGGTTCTTTGG |
| CCD8b-qRT | CAACTATGCCTTTTGGGTTAAAG | AAAGTCTCGGCCAAATCCT |
| DLK2C-qRT | CGATGTTGCCATATAGGTTGTGC | ACAAGGGAGCACACATGCAG |
| ZAS-qRT | TATGGAGGCCTTGCAAAACTTTGTC | CATTGTGTTTGCTAGTGATGATCTG |
| NSP2-qRT | TCAGCTGCTTCAACCACAGC | TGTTGGGACCCGTCTCCTC |
| CYCLOPS-qRT | GGTTTGGCTTGGTACAGCATCT | GGGAGGCAGGTCATCACAA |
| CCaMK-qRT | AGGCCAACAGCAAGTGATCT | CGCAGATTATCCAGCTCCTC |
| ABCB20-qRT | GAAATGCTTGATAGGGACACAC | TGAAACTCAGTTCTTCCCATGA |
| SYMRK-qRT | CCTGGCATAAAAGGGCAATA | GTGCTTTCGATGGACCTCAT |
| CERK1-qRT | TGGAATCGTGTACATCCCCG | CAAGTTGTGTGGAATCTTCAG |
| SLR1-qRT | GACGTCAACGAACGCTCAATT | CGGAGTCCAGTCGTCGATCT |
| PT11-qRT | ATATCCAAGGCCTCGTTCCT | CCGATCAGCTGGATCATGT |
| PT13-qRT | CAGGACGAGTATGGCCTCTT | TCGAGGACGAACCAACAGA |
| RLCK210 (KIN2)-qRT | CCTCATGGAGATGGACAAGAG | GATACCATCTCCTCCTCCAAAC |
| AM1-qRT | ACCGTGTGGGAGATGGAGTT | CCTGCAGCTCTTCCTCATCT |
| AM3-qRT | CTGTTGTTACATCTACGAATAAGGAGAAG | CAACTCTGGCCGGCAAGT |
| AM14-qRT | CCAACACCGTTGCAAGTACAATAC | GCACTTTGAAATTGGACTGTAAGAAA |
| PT2-qRT | GACGAGACCGCCCAAGAAG | TTTTCAGTCACTCACGTCGAGAC |
| PT6-qRT | CCGCCCCTGCAAACTGTA | CAACTGGCGGTTTCTTCGAT |
| OsUbiq-qRT | CATGGAGCTGCTGCTGTTCTAG | CAGACAACCATAGCTCCATTGG |
| CYCLOPHILIN2-qRT | AGCTCTCCTAGATCTGTGCTG | GCGATATCATAGAACGAGCGAC |
| IPS1-qRT | TTGGCAATTATTCGGTGGAT | ACCATTTCACCATCCTCTTTATG |
| GDPD2-qRT | GCCCAGTCATCTTCCATGATA | CCAATTCACTATCCGACCATCT |
| qPCR LjUbi | ATGCAGATCTTCGTCAAGACCTTG | ACCTCCCCTCAGACGAAG |
| qPCR LjPHR1A | CCGAATTGGAAGCATCCAAAGC | CTCGGAAGCTTGACTTTCGG |
| qPCR LjSYMRK | GAGGGTCAAAGGTGGATGA | GCGAACAATGGCGACCA |
| qPCR LjCCaMK | GGAGACAATGCAACTCTGTCTGA | CGGTGCTAGAGGGATCAATGA |
| qPCR LjCYCLOPS | GCTGGCAGATGAAAAAGAGC | GCGTGTTTGAGCACAACATT |
| qPCR LjD27 | GCCATCTCAATCGTTTATCAAG | GCTTCAGTGCTGGATCATC |
| qPCR LjCCD7 | GTATGGAGTGTTTAAGATGCCC | TAAAATGACTGCGTGGAAG |
| qPCR LjCCD8 | GGACACGCTTAGGAAATTCG | TCTGTCACAATGGGATGTGC |
| LjPHR1A genotyping | TTGGTTATAAAGGACCGCAAG | TTCCTAACTAAGCTTGCCCATAA |
| LORE1 P2 | | CCATGGCGGTTCCGTGAATCTTAGG |

**Table S2. Primers used for ChIP qPCR.**

Primers with name appended with “motif”, “left” and “right” were used for amplifying sequences flanking P1BS (GNATATNC), or 1000 bp left and 1000 bp right of P1BS motif respectively (-1000 bp and +1000 bp in Fig. S17). In case of *ECA1*pro, primers with name appended with “motif” represents P1BS-like (AMATATYC). In case of *CERK1*pro, primers with names appended with “GAGAmotif” and “CTTC motif” were used for amplifying sequences amplifying GGAGAGGA and TCCTCTTGTTCTTC elements respectively, while primer with names appended with “GAGAleft” and “CTTCright” were used for amplifying sequences 1000 bp left of GAGA and 1000 bp right of CTTC respectively.

| Primer name | Forward primer (5'→3') | Reverse primer (5'→3') |
|---|---|---|
| *CERK1*pro-ChIP-motif | TCGCAGTTTACAGTCGGAATC | TTGGATATACGGGCACACATTTA |
| *CERK1*pro-ChIP-left | GGCTGCTACATCACAAATTCAC | GGATGTGTTCGGCTGGTATT |
| *CERK1*pro-ChIP-right | AAGAACACAGAGTGAGCTGTAA | GGGAAGAAAGGGAGAAGAAGAG |
| *CCD7*pro-ChIP-motif | GGGCCTATAACTGCATATTCTCC | GTGCCCACGTAATTTGAAAGAG |
| *CCD7*pro-ChIP-left | CCTTCACTTGGCGTTACAGA | CAGACACTAAACAGCACTACGA |
| *CCD7*pro-ChIP-right | CATGCAGGTTCGTGGAGAC | TCACATTGCCCACCTTCTTC |
| *ZAS*pro-ChIP-motif | AGACACATGGATGCAGAGAAG | CGTGACGGATATTCCAAGATGA |
| *ZAS*pro-ChIP-left | CATTGGTGTTGCTGATGTTCTT | GCGGCCTACATTCTCAACTAT |
| *ZAS*pro-ChIP-right | GTCACATGGCATGCTACAAAC | AAATAACGGGTCCACCAATTTAAG |
| *SYMRK*pro-ChIP-motif | TGATTCCTCCCTTCCTCCTT | TTTCGTTCCGTGTCGTCATC |
| *SYMRK*pro-ChIP-left | ACAGTAACAAGGCTGAGTGTATC | AAGCAGCAATCCATCTACTCC |
| *SYMRK*pro-ChIP-right | TCTGCAGGACAACAACTTCA | GCAAATGGTAAAGAACAGCATCTA |
| *ECA1*pro-ChIP-motif | CGCACAGCAGAGCACAA | CCATTCTCCACTCTCCGTTTC |
| *ECA1*pro-ChIP-left | GAGATCGCATGGAACCGAAA | CTTCTCCTCTCTTCCGCATTG |
| *ECA1*pro-ChIP-right | TCGTGTAGGACAGACCTTGAT | AATCCTAGTCACCAGTCCTACC |
| *PT11*pro-ChIP-motif1 | CGAGAGGAGAATGACGAAATCA | GCTCTTCTCCCATATCCATCAG |
| *PT11*pro-ChIP-motif2 | TGATTGGCGATTCCTACCATAC | GCGTAGCGGTAAATCGATGA |
| *PT11*pro-ChIP-left | ATGCGCCACACGTAGTC | CCATGATCGTCTCTAGCATCTTC |
| *PT11*pro-ChIP-right | AGCGGTGAAGCAGCAAA | CTCTAGATAAGTGGGACCGTACA |
| *GDPD2*pro-ChIP-motif | TGCCTTTGGACCGGAATATC | AAGGAAGGAAGCGGGAATG |
| *GDPD2*pro-ChIP-left | TGTGCTCTCGTGATGAATCTG | CTGTTCCACGACGGGTTAAA |
| *GDPD2*pro-ChIP-right | TGAGCTGCTGTTCCGATTC | TGTCGATCGATTCGTATTTCCC |
| *NSP2*pro-ChIP-motif | GCATTACGGGAAGCAACAAAG | GCTGAACTGCTGAAGACTGA |
| *NSP2*pro-ChIP-left | CCATAGGTCGAGACTTGAGAG | CCCTTGGTACTTTAGAAATAGATGT |
| *NSP2*pro-ChIP-right | GCCTTGGCAACAAAGCTAAG | AGAAATGTGCCGAGAGAGATG |
| *CERK1*pro-ChIP-GAGAmotif | CAGTCCTGAACAGAGGACATAAG | GACTCCTCTCCAGACACTTCTA |
| *CERK1*pro-ChIP-CTTCmotif | GGAGTCAAGGTTAGTGGCTAAG | CTGTGTTCTTTGCTTACGGATG |
| *CERK1*pro-ChIP-GAGAleft | GTCTACCTCCACATGTCTCAAC | CCTAGGAAGAGGCCTAGATACA |
| *CERK1*pro-ChIP-CTTCright | TACCCGGCCAACAACATC | TGAGGAACAGCCCGTAGT |
| *RLCK210*pro-ChIP-motif | TTTGTGAATGAATTAGGTGCGT | GTGTTGTATGAGTATTGTCGAATGT |
| *RLCK210*pro-ChIP-left | TACTATCACACCACGCGTCTA | GAGATTAGTAGATGGTCCCTGTAATTT |
| *RLCK210*pro-ChIP-right | ACATGACCACCAGGCAAG | AATAAGACGGACGGTCAAACA |

**Table S3. Primers used for cloning.**

| Purpose | Name | Sequence |
|---|---|---|
| c*PHR2* cloning for p*35S*:c*PHR2* | MP503 | ATGAAGACTTTACGGGTCTCACACCATGGAGAGAATAAGCACCAATCAG C |
| | MP508 | ATGAAGACTTCAGAGGTCTCACCTTTCTGTCACCTGATTCT GAAACAAAAATTTAAGG |
| p*GDPD2* cloning for p*GDPD2*:GUS | MP595 | TTTGGTCTCAGCGGTGTTCATATATCTGATGTGACACGTC |
| | MP600 | TTTGGTCTCACAGATATATTCGGAGGATGTCCTAGCTG |
| p*GDPD2m* cloning for p*GDPD2m*:GUS | MP595 | TTTGGTCTCAGCGGTGTTCATATATCTGATGTGACACGTC |
| | MP596 | TTGGTCTCACGCGGACGGTCCAAAGGCACGC |
| | MP597 | TTGGTCTCACGCGAATGGAGGATAAACCATCCGATCCGC |
| | MP598 | TTGGTCTCAGGTCCGCGGAGGGGTGGGGATGCGTTC |
| | MP599 | TTGGTCTCAGACCCATTCCCGCTTCCTTC |
| | MP600 | TTTGGTCTCACAGATATATTCGGAGGATGTCCTAGCTG |
| p*PT11* cloning for p*PT11*:GUS | MP609 | TTGGTCTCTGCGGGGAGCAATAGACGAGGGATGCC |
| | MP614 | TTGGTCTCTCAGACTCCGATGATGCCGTCGATCG |
| p*PT11m* cloning for p*PT11m*:GUS | MP609 | TTGGTCTCTGCGGGGAGCAATAGACGAGGGATGCC |
| | MP610 | TTGGTCTCTTACGCGGAGGTAAATACATGAAAAATTAA AAGTTAGTTAGC |
| | MP611 | TTGGTCTCTCGTACACTGAACTACCCATTCACACC |
| | MP612 | TTGGTCTCTATTCGCGGAGGCAGATAATCATGATTG |
| | MP613 | TTGGTCTCTGAATACCAAAAACGACGCATTTCCGTCC |
| | MP614 | TTGGTCTCTCAGACTCCGATGATGCCGTCGATCG |
| p*CCD7* cloning for p*CCD7*:GUS | MP549 | TTTGGTCTCAGCGGGGGCGTGCACACTGCAAGCATC |
| | MP550 | TTTGGTCTCAATGATGTCTGCAAGGACCCAGAGCTCTAC |
| | MP551 | TTTGGTCTCATCATTCCTCTGTTCTTTCCACC |
| | MP552 | TTTGGTCTCACAGACTTTGGACTTGGCCTCCTTC |
| p*CCD7m* cloning for p*CCD7m*:GUS | MP549 | TTTGGTCTCAGCGGGGGCGTGCACACTGCAAGCATC |
| | MP591 | TTTGGTCTCATCCGCGTAAGTTATAGGCCCCGTTCGTTTTGG ATTTTGATGGCACATTTTTC |
| | MP592 | TTTGGTCTCACGGATCCGGGGAAAAATATTGAACTGGAATTAG |
| | MP552 | TTTGGTCTCACAGACTTTGGACTTGGCCTCCTTC |
| p*ZAS* cloning for p*ZAS*:GUS | MP545 | TTTGGTCTCAGCGGATATTTGGATGGTATGCAAAGCACATG |
| | MP546 | TTTGGTCTCAAGTTACGTACTCCCTCTGTTTCAC |
| | MP547 | TTTGGTCTCAAACTACGTACATATACCTAACGTAAC |
| | MP548 | TTTGGTCTCACAGATCTGCTAGTAAAAAAAGCCTAAATCC |
| p*ZASm* cloning for p*ZASm*:GUS | MP545 | TTTGGTCTCAGCGGATATTTGGATGGTATGCAAAGCACATG |
| | MP585 | TTTGGTCTCAGTACTCAGAAAAAAATTTCCGTCCCTTGTCC |
| | MP586 | TTTGGTCTCAGTACGCGGACAACGGGTCGTAGTCTTTAGTTATC |
| | MP587 | TTTGGTCTCATACGCGCAATAAAAAAGACGACAAAAAAAT ACATCATAAAAATCGATG |
| | MP588 | TTTGGTCTCACGTAGCATGGTTTTTTTCTTTTCTTTCCAG |
| | MP589 | TTTGGTCTCAGAAGTCACGGAACCATCTTGGTG |
| | MP590 | TTTGGTCTCACTTCGCGGACAAGATGACAAATGGAATTTCATCAC |
| | MP548 | TTTGGTCTCACAGATCTGCTAGTAAAAAAAGCCTAAATCC |

**Table S4. Plasmid construction by Golden Gate cloning (Level I, II and III).** The Golden Gate toolbox was previously described[60].

| Purpose | Name | Description |
|---|---|---|
| **Golden Gate level I (LI) elements** | | |
| | LI c*PHR2* | PCR amplification of *OsPHR2* coding sequence from Nipponbare cDNA with MP503 + MP508 and assembly by BpiI cut ligation into LI pUC57 plasmid (BB03). |
| | LI C-D *GUS* | ref. 81 |
| **Golden Gate level II (LII) plasmids** | | |
| | LII F 3-4 p*35S:cPHR2* | Assembled by BsaI cut ligation from: LI A-C p*35S* (G009) + LI dy B-C (BB6) + LI c*PHR2* + LI D-E c-Myc (G070) + LI E-F 35S-T (G059) + LI dy F-G (BB09) + LII R 3-4 (BB24) |
| | LIIc F 1-2 p*Ubi:mCherry* | Assembled by BsaI cut ligation from: LI A-B p*Ubi* (G007) + LI B-C (BB06) dy + LI C-D *mCherry* (G023) + LI D-E (BB08) dy + LI E-F 35S-T (G059) + LI F-G dy (BB09) + LIIc F 1-2 (BB30) |
| | LIIc R 5-6 p*35S*:*mCherry* | Assembled by BsaI cut ligation from: LI A-B p*35S* (G009) + LI B-C (BB06) dy + LI C-D *mCherry* (G023) + LI D-E (BB08) dy + LI E-F 35S-T (G059) + LI F-G dy (BB09) + LIIc R 5-6 (BB30) |
| | LIIc F 3-4 pOI:*GUS* | Assembled by BsaI cut ligation from: LI A-B Esp3I-*lacZ* dy (G082) + LI B-C dy (BB06) + LI C-D *GUS* + LI D-E dy (BB08) + LI nos-T (G006) + LI F-G dy (BB09) + LIIc F 3-4 (BB33) |
| **Golden Gate level III (LIII) plasmids for plant transformation** | | |
| *Overexpression of OsPHR2 in N. benthamiana* leaves | LIIIβ F A-B p*35S:cPHR2* | Assembled by BpiI cut ligation from: LII dy 1-2 (BB63) + LII dy 2-3 (BB39) + LII F 3-4 p*35S:cPHR2* + LII dy 4-5 ins (BB44) + LIIc R 5-6 p*35S*:*mCherry* + LIIIβ F A-B (BB53) |
| Esp3I compatible destination backbone for Localization of promoter activity | Esp3I cut ligation compatible backbone: LIIIβ fin p*Ubi:mCherry*_pOI:*GUS* Esp3I | Assembled by BpiI cut ligation from: LIIc F 1-2 p*Ubi:mCherry* + LII 2-3 ins (BB43) + LIIc F 3-4 pOI:*GUS* + LII dy 4-6 (BB41) + LIIIβ fin (BB52) |
| BsaI compatible destination backbone for Localization of promoter activity | BsaI cut ligation compatible backbone: LIIIβ fin p*Ubi:mCherry*_pOI:*GUS* BsaI | Assembled by Esp3I cut ligation from: LIIIβ fin p*Ubi:mCherry*_pOI:*GUS* Esp3I + LI A-B Esp3I-ccdB dy (G084) |
| Transactivation of p*GDPD2*:GUS in *N. benthamiana* leaves | LIIIβ fin p*Ubi*:*mCherry*_p*GDPD*:*GUS* | Assembled by BsaI cut ligation from: LIIIβ fin p*Ubi:mCherry*_pOI:*GUS* BsaI + PCR amplicon MP595 + MP600 amplified from Nipponbare genomic DNA |
| Transactivation of p*GDPD2m*:GUS in *N. benthamiana* leaves | LIIIβ fin p*Ubi*:*mCherry*_p*GDPDm*:*GUS* | Assembled by BsaI cut ligation from: LIIIβ fin p*Ubi:mCherry*_pOI:*GUS* BsaI + PCR amplicons MP595 + MP596, MP597 + MP598, MP599 + MP600 |

| | | |
|---|---|---|
| | | amplified from LIIIβ fin p*Ubi*:*mCherry*_p*GDPD*:*GUS* |
| Transactivation of p*PT11*:GUS in *N. benthamiana* leaves | LIIIβ fin p*Ubi*:*mCherry*_p*PT11*:*GUS* | Assembled by BsaI cut ligation from: LIIIβ fin p*Ubi:mCherry*_pOI*:GUS* BsaI + PCR amplicon MP609 + MP614 amplified from Nipponbare genomic DNA |
| Transactivation of p*PT11m*:GUS in *N. benthamiana* leaves | LIIIβ fin p*Ubi*:*mCherry*_p*PT11m*:*GUS* | Assembled by BsaI cut ligation from: LIIIβ fin p*Ubi:mCherry*_pOI*:GUS* BsaI + PCR amplicons MP609 + MP610, MP611 + MP612, MP613 + MP614 amplified from LIIIβ fin p*Ubi*:*mCherry*_p*PT11*:*GUS* |
| Transactivation of p*CCD7*:GUS in *N. benthamiana* leaves | LIIIβ fin p*Ubi*:*mCherry*_p*CCD7*:*GUS* | Assembled by BsaI cut ligation from: LIIIβ fin p*Ubi:mCherry*_pOI*:GUS* BsaI + PCR amplicons MP549 + MP550, MP551 + MP552 amplified from Nipponbare genomic DNA |
| Transactivation of p*CCD7m*:GUS in *N. benthamiana* leaves | LIIIβ fin p*Ubi*:*mCherry*_p*CCD7m*:*GUS* | Assembled by BsaI cut ligation from: LIIIβ fin p*Ubi:mCherry*_pOI*:GUS* BsaI + PCR amplicons MP549 + MP591, MP592 + MP552 amplified from LIIIβ fin p*Ubi*:*mCherry*_p*CCD7*:*GUS* |
| Transactivation of p*ZAS*:GUS in *N. benthamiana* leaves | LIIIβ fin p*Ubi*:*mCherry*_p*ZAS*:*GUS* | Assembled by BsaI cut ligation from: LIIIβ fin p*Ubi:mCherry*_pOI*:GUS* BsaI + PCR amplicons MP545 + MP546, MP547 + MP548 amplified from Nipponbare genomic DNA |
| Transactivation of p*ZASm*:GUS in *N. benthamiana* leaves | LIIIβ fin p*Ubi*:*mCherry*_p*ZASm*:*GUS* | Assembled by BsaI cut ligation from: LIIIβ fin p*Ubi:mCherry*_pOI*:GUS* BsaI + PCR amplicons MP545 + MP585, MP586 + MP587, MP588 + MP589, MP590 + MP548 amplified from LIIIβ fin p*Ubi*:*mCherry*_p*ZAS*:*GUS* |
